# Mendelian Randomization Analysis Using Multiple Biomarkers of an Underlying Common Exposure

**DOI:** 10.1101/2021.02.05.429979

**Authors:** Jin Jin, Guanghao Qi, Zhi Yu, Nilanjan Chatterjee

## Abstract

Mendelian Randomization (MR) analysis is increasingly popular for testing the causal effect of exposures on disease outcomes using data from genome-wide association studies. In some settings, the underlying exposure, such as systematic inflammation, may not be directly observable, but measurements can be available on multiple biomarkers or other types of traits that are coregulated by the exposure. We propose a method for MR analysis on latent exposures (MRLE), which tests the significance for, and the direction of, the effect of a latent exposure by leveraging information from multiple related traits. The method is developed by constructing a set of estimating functions based on the second-order moments of GWAS summary association statistics for the observable traits, under a structural equation model where genetic variants are assumed to have indirect effects through the latent exposure and potentially direct effects on the traits. Simulation studies show that MRLE has well-controlled type I error rates and enhanced power compared to single-trait MR tests under various types of pleiotropy. Applications of MRLE using genetic association statistics across five inflammatory biomarkers (CRP, IL-6, IL-8, TNF-α, and MCP-1) provide evidence for potential causal effects of inflammation on increasing the risk of coronary artery disease, colorectal cancer, and rheumatoid arthritis, while standard MR analysis for individual biomarkers fails to detect consistent evidence for such effects.

## 1. Introduction

In the last two decades, Mendelian Randomization (MR) analysis has become increasingly popular for investigating the causal effects of risk factors and biomarkers on disease outcomes when randomized controlled trials are unavailable (Davey Smith and Ebrahim, 2003; Lawlor *and others*, 2008; Emdin *and others*, 2017). MR analysis is a type of instrumental variable (IV) analysis that uses genetic variants (SNPs) as “instruments” aiming for an unbiased estimate of the underlying causal effect when there exist potential confounders. While early MR analysis focused on using individual genetic variants of known functional consequence as instruments (Greenland, 2000; Abifadel *and others*, 2003; Cohen *and others*, 2006; Lawlor *and others*, 2008), the era of genomewide association studies (GWAS) has led to the rise of powerful MR analysis methods based on multiple SNPs (Pierce *and others*, 2011; Pierce and Burgess, 2013; Evans and Davey Smith, 2015; Zheng *and others*, 2017a). In particular, with the rapid development of large publicly available GWAS, the majority of MR research has been transferred from working with individual-level data to two-sample MR analysis with summary-level data (Bautista *and others*, 2006; Burgess and Thompson, 2015; Qi and Chatterjee, 2021).

Many recent methods for MR analysis have focused on improving robustness to the presence of pleiotropic association by which genetic instruments can affect the outcome of interest independent of the exposure and thus leading to violation of key assumptions (Bowden *and others*, 2015, 2016; Hartwig *and others*, 2017; Verbanck *and others*, 2018; Zhu *and others*, 2018; Qi and Chatterjee, 2019; Morrison *and others*, 2020; Shapland *and others*, 2022; Zhao *and others*, 2020; Cinelli *and others*, 2022). In this study, we consider a novel setting of pleiotropic association, where genetic variants might be associated with multiple observed traits through an underlying latent exposure which may have a causal effect on the outcome. An example of the relevant issues has recently been illustrated in a study of the causal relationship between blood pressure and multiple kidney function biomarkers (Yu *and others*, 2020). The analysis showed that standard MR analysis methods using an individual kidney function biomarker, such as the estimated glomerular filtration rate determined based on the serum creatinine (eGFRcr), did not detect a causal effect of kidney function on blood pressure. Instead, when the analysis was restricted to instruments that showed association across multiple biomarkers and thus were likely to be related to the underlying kidney function, it showed clear evidence of causal effects. Examples of groups of traits that may be governed by an underlying common exposure appear in many settings, including but not limited to, inflammatory biomarkers (Brenner *and others*, 2014; Qian *and others*, 2019), groups of metabolites related to dietary and lifestyle exposures (Gu *and others*, 2018; Oluwagbemigun *and others*, 2020), and measurement instruments for evaluations of underlying mental health conditions (Tsanas *and others*, 2017; Black *and others*, 2019).

We propose a novel method for conducting MR analysis on latent exposures (MRLE), which allows testing for the statistical significance and direction of the effect of an unobservable latent exposure by leveraging information from multiple related observable traits. The method uses GWAS summary-level association statistics of the observable traits and the outcome on a set of “strictly selected” SNPs (IVs) that are associated with at least two of the traits at specified thresholds of significance. We use an underlying structural equation model to describe causal paths between the SNPs, the latent exposure, the traits co-regulated by the exposure, and the outcome. We then construct a series of estimating functions by equating the second-order sample moments of the summary-level association statistics with the corresponding theoretical moments and propose inference for identifiable parameters based on the Generalized Method of Moments theory (Hansen, 1982; Newey and McFadden, 1994; Hall *and others*, 2005).

We show by simulation that the proposed MRLE test has a well-controlled type I error rate, increased power, and a higher probability of correctly identifying the direction of the causal effect compared to single-trait MR analysis, and is robust to the presence of various types of pleiotropy across the latent exposure, the observable traits, and the outcome. We applied MRLE to test the effect of chronic inflammation on rheumatoid arthritis, coronary artery disease, and cancers including colorectal cancer, prostate cancer and endometrial cancer, using up to five inflammatory biomarkers including c-reactive protein (CRP), interleukin 6 (IL-6), interleukin 8 (IL-8), tumor necrosis factor alpha (TNF-α), and monocyte chemoattractant protein-1 (MCP-1). These analyses reveal that while MR analyses based on single biomarkers often fail to detect any consistent patterns of causal effects, the proposed MRLE method detects such evidence for a number of diseases including RA, CAD, and CRC.

The rest of the paper is organized as follows. In Section 2, we introduce the model setup and method development for MRLE. We investigate the performance of MRLE in a variety of simulated data scenarios in Section 3 and through an application of testing the causal effect of chronic inflammation on multiple diseases in Section 4. Further discussion and future directions are presented in Section 5.

## 2. Methods

### 2.1 Problem setting and model setup

Let *X* denote an exposure variable of interest for which we want to examine its causal effect *θ* on an outcome *Y*. We assume that measurements on *X* are not directly available, and instead one can observe a set of biomarkers, or other types of traits, {*B_k_, k* = 1,…, *K*}, *K* ⩾ 2, that are co-regulated by *X* with known directions. We call a biomarker “valid” for such analysis on latent exposure *X* if it meets the following two conditions: (A) *X* affects the biomarker, and (B) conditional on *X*, the biomarker does not have an effect on the outcome itself (“pure surrogate”). Suppose we select *M* independent SNPs *G_j_, j* = 1,2,…, *M*, that are associated with one or more of the *B_k_*s, as instrumental variables (IVs) for the MR analysis. We assume that a nonzero proportion of the selected IVs are valid IVs for *X*, i.e., they are directly associated with *X*. Details of the IV selection strategy will be discussed later in Section 2.3. Following we will assume that the outcome *Y* is continuous and develop our method in the linear structural equation modeling framework. Note that we show method development for a continuous outcome *Y* for simplicity, but the testing result is invariant with respect to the scaling of the effect size, and thus the method can be applied to discrete (e.g., binary/categorical) outcomes without further modification (see Appendix B of the Supplementary Materials for details). We further assume without loss of generality that the SNPs *G_j_, j* = 1,2, …,*M*, the biomarkers *B_k_, k* = 1, 2,…, *K*, the latent exposure *X*, and the outcome *Y* are all standardized to have unit variance, so that the effect *θ* is on the standardized scale. Again, such an assumption is only made to simplify notations in our derivations: even if the data is not standardized, the method still works since we only care about the existence and direction of the causal effect but not the effect size.

Figure 1 describes the assumed causal paths between the SNPs (*G_j_*s), the latent exposure of interest (*X*), the biomarkers co-regulated by the latent exposure (*B_k_*s), and the outcome (*Y*), assuming a total of *K* = 3 biomarkers. Here we assume an “outcome model” of the form

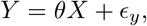

where *ϵ_y_* denotes a mean-zero error term with variance 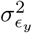. Here *ϵ_y_* can be correlated with *X* due to unobserved confounding for the effect of *X* on *Y*, but we assume the confounders are not associated with the selected IVs. Our goal is to conduct a hypothesis test with null hypothesis *H*_0_: *θ* = 0 and alternative hypothesis *H_a_*: *θ* ≠ 0, also to infer the sign of *θ* when the null hypothesis is rejected.

**Fig. 1:**
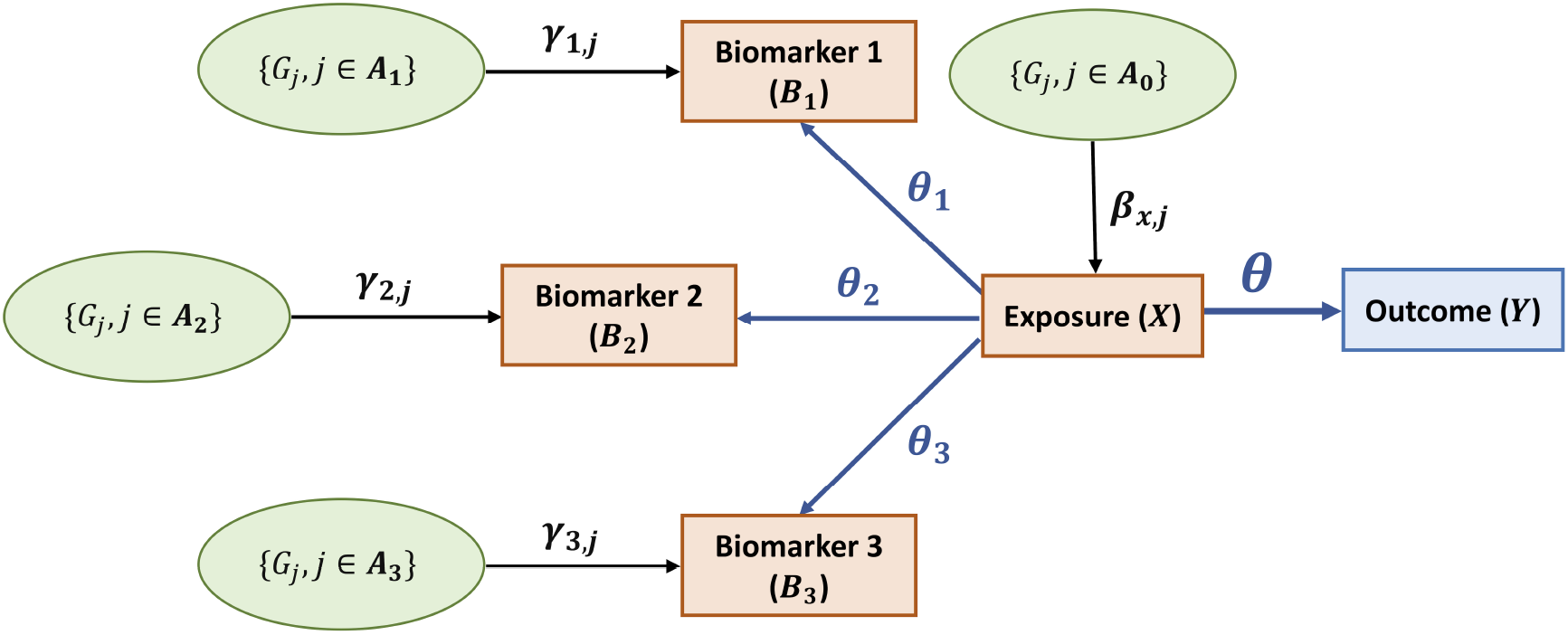
Causal paths between the SNPs (*G_j_*s), the latent exposure of interest (*X*), the biomarkers co-regulated by the latent exposure (*B_k_*s), and the outcome (*Y*). *A*_0_ represents the set of indexes for SNPs that are directly associated with *X* with effect sizes *β_x,j_*s, and Ak represents the set of indexes for SNPs that are not associated with *X* but directly associated with *B_k_* with effect sizes *γ_k,j_*s, *k* = 1,…, *K*. *A*_0_ and *A_k_*s may have overlaps with each other, but the effects of one SNP on different traits or the outcome are assumed independent, although we will show validity of the proposed MRLE test on various correlated pleiotropy settings by simulations. The number of observable traits is set to *K* = 3 for illustration.

We next assume an “exposure model” of the form

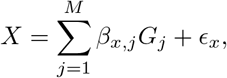

where *β_x,j_, j* = 1, 2,…, *M* denote additive effects of SNPs on *X*, and *ϵ_x_* is a mean-zero error term with variance 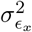. We further assume that the effect sizes, *β_x,j_, j* = 1,2,…, *M*, can be modelled as i.i.d. mean-zero random variables with variance 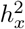.

Finally, we assume a set of “biomarker models” in the form

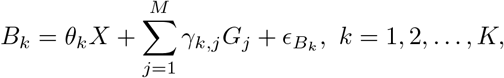

where *θ_k_* denotes the effect of exposure *X* on biomarker *B_k_, γ_k,j_, j* = 1,…, *M* denote the direct effects of the SNPs on *B_k_* that are not medicated through *X*, and *ϵ_B_k__* denotes the mean-zero residual error term associated with *B_k_* with a variance 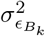. We further assume that for each *k, γ_k,j_, j* = 1,2…, *M* are i.i.d. mean-zero random variables with a variance 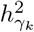 that are uncorrelated with association coefficients for SNPs associated with *X*, i.e., *β_x,j_, j* = 1,…, *M*, in the “exposure” model.

Assume we have summary-level association statistics for biomarker 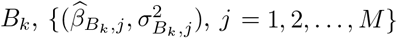, and those for the outcome 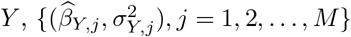, from separate GWAS, where 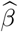 and *σ*^2^ with specific subscript denote the corresponding estimated association coefficient and its estimated standard error, respectively. We allow the GWAS for *B_k_*s to have potentially overlapping samples but assume that GWAS for *Y* is independent of GWAS for *B_k_*s to avoid introducing bias caused by sample overlap in two-sample MR (Burgess, Davies, and Thompson, 2016).

It has been common practice to test for causal effect of an underlying exposure, such as chronic inflammation, using a surrogate biomarker such as CRP, based on a standard IVW estimator, 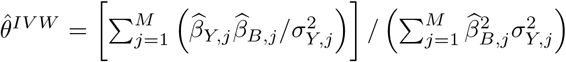. Under the above structural equation model, we can derive the asymptotic bias of the standard IVW estimator in the form of 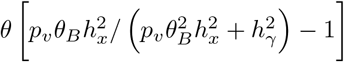, as GWAS sample sizes go to infinity. Here *p_v_* ∈ [0, 1] denotes the proportion of IVs selected for the biomarker that are valid IVs for the latent exposure, i.e., the rest of the SNPs are directly associated with the biomarker but not directly associated with the latent exposure (see Appendix A of the Supplementary Materials for mathematical details). Thus, under the above model, while a standard test based on an individual biomarker is expected to be valid with a well-controlled type I error rate given a zero asymptotic bias when *θ* = 0, it can lose major power due to attenuation of the underlying causal effect when 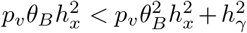. Furthermore, as we will show later, in the presence of more complex pleiotropy settings, single biomarker-based tests for causal effects can easily become invalid with inflated type I error rates.

### 2.2 Cross-biomarker MR analysis using generalized method of moments

We propose statistical inference based on the method of moments which only requires moments of the summary-level association statistics and no additional distributional assumptions for model parameters. Let 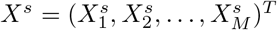 denote the observed GWAS summary-level data for biomarkers and outcome, where 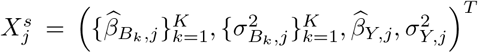 for SNP *j* = 1, 2,…, *M*. We consider the second-order moments of 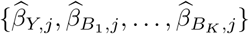, which, based on our model described in the previous section, can be derived as

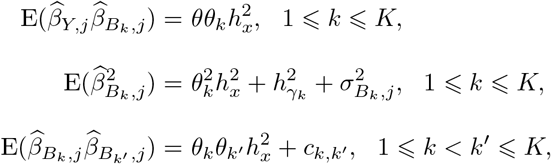

where *c_k,k′_* denotes the covariance between 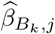 and 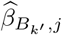, which could arise due to sample overlap between GWAS of correlated biomarkers, and can be estimated from summary-level association statistics using techniques such as bivariate LD score regression (Turley *and others*, 2018; Qi and Chatterjee, 2018). Detailed derivations are provided in Appendix B of the Supplementary Materials.

We note that the per-SNP heritability of the latent exposure 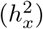 always shows up alongside the terms 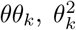, or *θ_k_θ_k′_*, *k* = 1, 2,…, *K*, which makes it impossible to infer 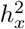 separately from *θ* and *θ_k_*s. Such an identifiability issue is actually the reason we cannot estimate the causal effect (θ) of *X* on *Y* if not fixing 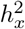. To avoid such an issue, we reparameterize the model with *μ* = *θh_x_, μ_k_* = *θ_k_h_x_, k* = 1, 2,…, *K*, and denote the vector of model parameters by 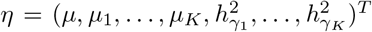. Conducting inference on *θ* is then equivalent to conducting inference on *μ*: not only the hypothesis test 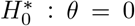 versus 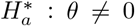 is equivalent to 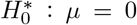 versus 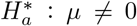, but also the sign of *θ* is the same as the sign of *μ* = *θh_x_*, given that *h_x_* (i.e., standard deviation of *β_x,j_*s) is positive. Due to the existence of the terms 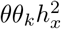, *k* = 1, 2,…, *K*, the identifiability of the sign of *θ* requires that the signs of *θ_k_*s, i.e., the associations between *B_k_*s and *X*, are known. Such an assumption is implicit in our proposed MRLE method, which is reasonable because the directions of the effect of latent exposure on biomarkers are typically known and available in the literature. One example is that for kidney function, it is known that a higher level of estimated glomerular filtration rate (eGFR) based on either serum creatinine (eGFRcr) or cystatin C (eGFRcys), a commonly used kidney function biomarker, indicates better kidney function, while a higher level of blood urea nitrogen (BUN), a complementary kidney function biomarker, indicates weaker kidney function (Levey *and others*, 2009; Yu *and others*, 2020). Another example is on chronic inflammation, the latent exposure we will discuss in Section 4. Chronic inflammation has multiple commonly seen biomarkers, such as CRP, IL-6, IL-8, TNF-α, and MCP-1, that are all known to be positively associated with chronic inflammation (Sproston and Ashworth, 2018). But even if the signs of some *θ_k_*s are unknown, the MRLE test is still valid, where a rejection of *H*_0_ suggests a causal effect of the latent exposure on the outcome, but we cannot determine its direction.

By equating the second-order moments of the summary-level association coefficients with the corresponding sample moments, we obtain our estimating functions, 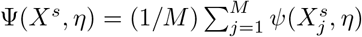, where 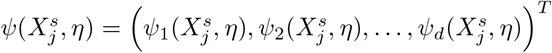, and estimating equations Ψ(*X^s^, η*) = 0, with

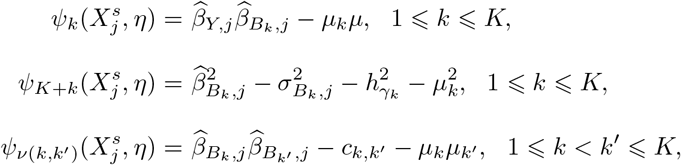

where 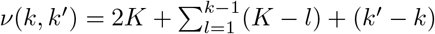 and *d* = *K*(*K* + 3)/2.

We observe that there are *d* = *K*(*K*+3)/2 equations and *p* = 2*K* +1 unknown parameters. To solve the estimating equations, we need *d* ⩾ *p,* i.e., *K* ⩾ 2 biomarkers are required. When *K* = 2, the number of estimating equations equals the number of unknown parameters, and we can solve the equations directly. When *K* ⩾ 3, there are more equations than unknown parameters, the problem becomes over-identified and we may not be able to solve the exact equations. To deal with this issue, we consider the generalized method of moments (GMM) and define the following GMM estimator, 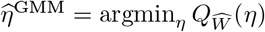, which is the global minimizer of the objective function 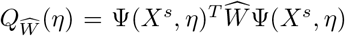 where 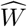 is a positive semi-definite weighting matrix which is typically specified based on *X^s^*. Here we consider the optimal choice of 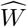 with a minimum asymptotic covariance, 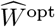, which is the inverse of 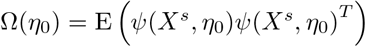, i.e., the covariance matrix of the estimating equations based on true parameter values *η*_0_. The closed form expression of Ω(*η*_0_) can be easily derived, but since *η*_0_ is unknown, in applications we conduct inference using an iterative algorithm. Specifically, we implement a two-step GMM algorithm where in step one we obtain an initial estimate of *η* with a simple choice of 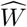, such as the identity matrix, and in step two we estimate *η* iteratively by setting 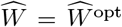 based on the estimate from the previous iteration until a pre-specified convergence criterion is met. Derivation of 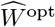 and detailed algorithm are provided in Appendix C of the Supplementary Materials. Inference on the causal effect *θ* can then be conducted based on 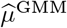, i.e., the first element of 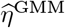. We obtain 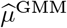 and an estimate of its standard error based on the GMM theory, then perform a Wald test on *H*_0_: *μ* = 0 versus *H_a_*: *μ* ≠ 0. Properties of 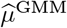 and computational details are summarized in Supporting Information Web Appendices B and C.

### 2.3 Selection of genetic IVs

A key issue in simultaneously analyzing multiple biomarkers is how to select instruments that may be associated with the underlying common exposure. As noted earlier, many SNPs that are associated with individual biomarkers may represent genetic variations that are unrelated to the latent exposure, and using these SNPs as IVs may lead to power loss. Furthermore, if there are numerous biomarkers, the chance of incorporating exposure-independent pleiotropic association of SNPs with some of the biomarkers also increases, potentially leading to bias.

To alleviate these issues, we propose a more strict IV selection criterion, where we select SNPs that are associated with at least two of the biomarkers co-regulated by the latent exposure. Intuitively, SNPs associated with multiple biomarkers are more likely to be directly related to the underlying common exposure and less likely to suffer from pleiotropic effects due to any shared genetic background between individual biomarkers and the outcome. However, the use of GWAS significance threshold (5 × 10^-8^) can be overly strict for this IV selection strategy, especially when GWAS sample sizes for some biomarkers are relatively small. For example, in our data analysis which will be described in Section 4 in detail, while GWAS for CRP was large (*N* = 320041), those for the other biomarkers were relatively small (*N* = 3454 ~ 8394). To ensure that a minimum number of IVs are selected, a more liberal threshold can be used depending on the GWAS sample size. We show by simulation that such a strategy of selecting IVs that affect multiple biomarkers can effectively reduce pleiotropic bias, leading to a higher power of detecting causal effect of the latent exposure (Sections 3.2, 3.3, 3.4, and 3.5) and avoiding highly inflated type I error rates (Section 3.3), under various types of pleiotropy settings.

## 3. Simulation Studies

### 3.1 Simulating individual-level versus summary-level data

In the following sections, we will illustrate the performance of the proposed MRLE method by simulating genome-wide studies under a variety of scenarios. Conducting large-scale, genomewide simulations with data being generated on the individual level is highly computationally intensive. We, therefore, considered simplifying the simulation procedure by directly simulating GWAS summary statistics, which is much faster to implement.

We first conducted a pilot simulation study to show the consistency of the results between simulating on the individual level and simulating on the summary level, which is expected from the theory. Detailed simulation settings are summarized in Appendix D.1 of the Supplementary Materials. For both individual-level and summary-level simulations, we assumed there are unmeasured confounders between the exposure, the biomarkers, and the outcome, but the selected genetic variants satisfy the key “instrumental variable” assumption that they are not related to the confounders. Specifically, we assume a correlation 0.3 between the residual term *ϵ_y_* in the “outcome model”, *Y* = *θX* + *ϵ_y_*, and the residual term *ϵ_x_* in the “exposure model”, 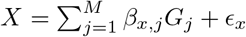, and similarly, a correlation 0.3 between *ϵ_y_* or *ϵ_x_* and the residual term *ϵ_B_k__*s in the “biomarker models”, 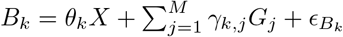, *k* = 1, 2,…, *K*.

We set the GWAS sample size to *N* = 6 × 10^4^ for the outcome and each of the *K* = 6 biomarkers that are co-regulated by the latent exposure. For individual-level simulations, we collected genotype data for *N* unrelated individuals that were randomly selected from the UK Biobank samples. We first conducted LD pruning to select M=126,627 relatively independent SNPs to be included in our analysis. Instead of assuming all SNPs to be causal as in the “exposure” and “biomarker” model, we considered a more realistic setting, where only a small proportion (1%) of the SNPs are causal with non-zero effect on the biomarker or the exposure. We generated the true effects of the SNPs and simulated data for the exposure, the biomarkers, and the outcome for the *N* individuals based on our assumed model. We then conducted GWAS analysis on the outcome and each biomarker using one-SNP-at-a-time regressions. The various tests were then applied to the GWAS summary data. Summary-level simulations were conducted under the exact same simulated data scenario as in individual-level simulations, except that we directly simulated GWAS summary data according to the derived distribution of the summary-level association statistics (Appendix D of the Supplementary Materials).

The consistency of the hypothesis testing results between individual-level and summary-level simulations is illustrated in an example data scenario. We observe from Figure 2 that the two types of simulations have similar rejection rates. As expected, both MRLE and IVW provide valid tests for the causal effect of the latent exposure, with type I error rates well controlled at approximately *α* = 0.05 in the presence of confounding effects. The computation time required for completing 1000 individual-level simulations with a sample size of *N* = 6 × 10^4^ is approximately 142 hours. On the other hand, 1000 summary-level simulations under the same data scenario were completed within 2.13 hours. Considering that we will conduct large-scale, genome-wide simulations under a large number of data scenarios, summary-level simulation is much more computationally feasible. In our main simulation study (Sections 3.2 - 3.5), we thus directly simulated GWAS summary data to reduce the computational burden.

**Fig. 2:**
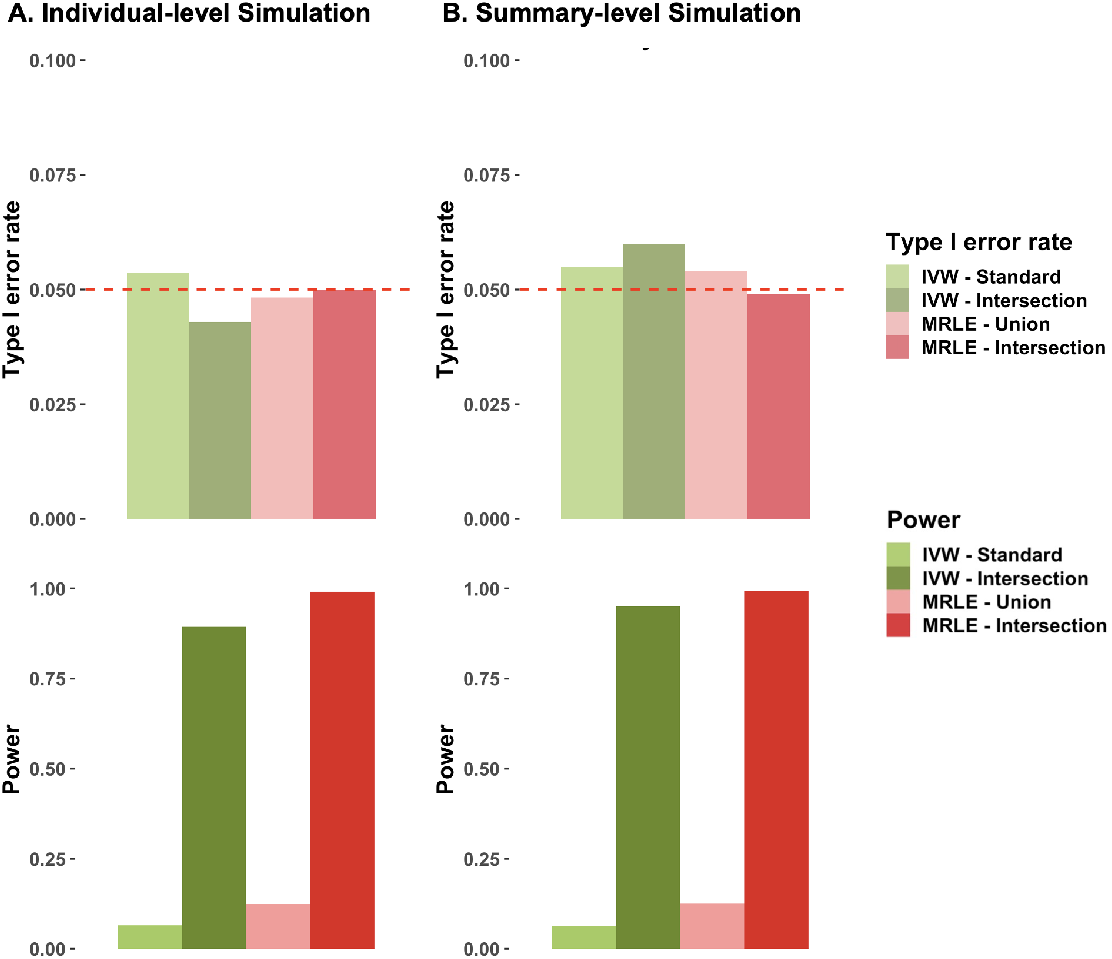
Results from the pilot simulation study showing consistency of the results between individual-level simulation **(A)** and summary-level simulation **(B)**. Results were summarized from 1000 simulations assuming a total of *K* = 6 biomarkers, with a GWAS sample size of *N* = 6 × 10^4^ for the outcome and all biomarkers. The total heritability of each biomarker 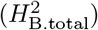 and the heritability of each biomarker explained by the latent exposure 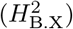 were both set to 0.2. IVs are defined as either the SNPs associated with at least one biomarker (“IVW-Standard” and “MRLE-Union”, *α* = 5 × 10^-8^) or the SNPs associated with at least two biomarkers (“IVW-Intersection” and “MRLE-Intersection”, *α* = 5 × 10^-6^). In each subfigure, the upper panel shows the empirical type I error rate under *θ* = 0, and the lower panel shows the empirical power under *θ* = 0.1.

### 3.2 Simulations assuming no pleiotropy

Suppose there are *M* = 2 × 10^5^ independent common SNPs across the whole genome and the summary-level data are available for *K* = 4, 6 or 8 biomarkers representing an underlying latent exposure. As mentioned in Section, instead of assuming all SNPs to have non-zero effect on the trait (exposure or biomarker), we considered a more realistic setting where only a small proportion of the SNPs are causal, i.e., having non-zero effect on each trait. Specifically, we assumed that a random subset of *M_x_* = *π_x_M* SNPs were associated with *X*, with effect sizes *β_x,j_*s generated from 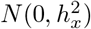; similarly, a random subset of *M_B_k__* = *π_B_k__M* SNPs were associated with *B_k_*, with effect sizes *γ_k,j_*s generated from 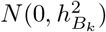, *k* = 1, 2,…, *K*. We set the proportion of causal SNPs to *π_x_* = *π_B_k__* = 1%, and *θ_k_* to 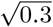, *k* = 1, 2,…, *K*, so that *X* explains 30% of variability of each of the biomarkers. We further set the total heritability of each biomarker 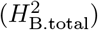 to 0.2 or 0.3, and the proportion explained by the association with *X* to 0.2 or 0.3, which leads to a total heritability of each biomarker that is explained by *X* 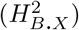 between 0.04 and 0.09.

As mentioned earlier, we directly simulated GWAS summary-level association statistics to avoid having to simulate large-scale individual-level data (Qi and Chatterjee, 2018, see Appendix D.2 of the Supplementary Materials for details). For simplicity, we assumed equal sample sizes across all GWAS and all SNPs, which were set equal to 6 × 10^4^, 8 × 10^4^, or 10^5^. We also set the overlapping GWAS sample size between any two biomarkers to *N_B_k_,B_l__* = *N*, i.e., the GWAS summary data for all Biomarkers were obtained from the same set of individuals, and between-biomarker correlation to *cov*(*B_k_, B_l_*) = 0.3, 1 ⩽ *k < l* ⩽ *K*. We also set *N_B_k__, Y* = 0 for *k* = 1,…, *K* based on the requirement of no sample overlap between the biomarkers and the outcome.

After generating summary-level association statistics, we selected IVs using either one of the two strategies discussed in Section 2.3, i.e., we selected either (1) the union of the SNPs that reached genome-wide significance threshold (5 × 10^-8^) for any single *B_k_*, the corresponding test was denoted by “MRLE-Union”; or (2) SNPs that reached a more liberal significance threshold, 5 × 10^-6^, for at least two biomarkers, the corresponding test was denoted by “MRLE-Intersection”. As a comparison, we also applied the fixed-effect IVW tests based on SNPs associated with each individual biomarker. Similiar to MRLE, we also considered two IV selection strategies, where for each single-biomarker IVW test, we selected either (1) SNPs that reached genome-wide significance threshold (5 × 10^-8^) for that biomarker only (“IVW-Standard”); or (2) SNPs that reached significance level 5× 10^-6^ for that biomarker and at least one other (“IVW-Intersection”). Since multiple IVW tests were conducted, an adjusted significance level, *α*_1_ = 1 – (1 – *α*_0_)^1/K^ was used for the test on each biomarker to control family-wise error rate (FWER) at *α*_0_. We assessed type I error control of the various methods at *θ* = 0 and power at *θ* = 0.1.

Under the no pleiotropy assumption, type I error rates seem to be well controlled at approximately *α*_0_ = 0.05 in all simulated settings (Figures 3, S2, and S3). We observe that overall, as GWAS sample size (N) increases (which leads to increased number of IVs), the power of both IVW and MRLE tests increases (Figure 3). The power of the tests also increases as the total heritability of the biomarkers 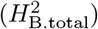 and the proportion of this heritability explained by the exposure 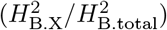 increase. Compared to the naive IV selection strategy of choosing all SNPs associated with any biomarker, the proposed, more strict criterion of choosing SNPs associated with at least two biomarkers yields a higher power and a higher probability of correctly identifying the direction of the effect for both tests (Figures 3, S2, S3, and S4(B), MRLE-Intersection versus MRLE-Union, IVW-Intersection versus IVW-Standard). Using either of the two IV selection strategies, MRLE provides a substantially higher power and a higher probability of correctly identifying the causal direction compared to the IVW test under the same level of type I error control. We also conducted simulation studies under the same settings but assuming a total of *K* = 4 or 8 biomarkers (Figures S2, S3, and S4(A,C)). Overall, the results are similar to those presented in Figures 3 and S4(B), but using a larger number of biomarkers tends to give a higher power and a higher chance of correctly identifying the causal direction.

**Fig. 3:**
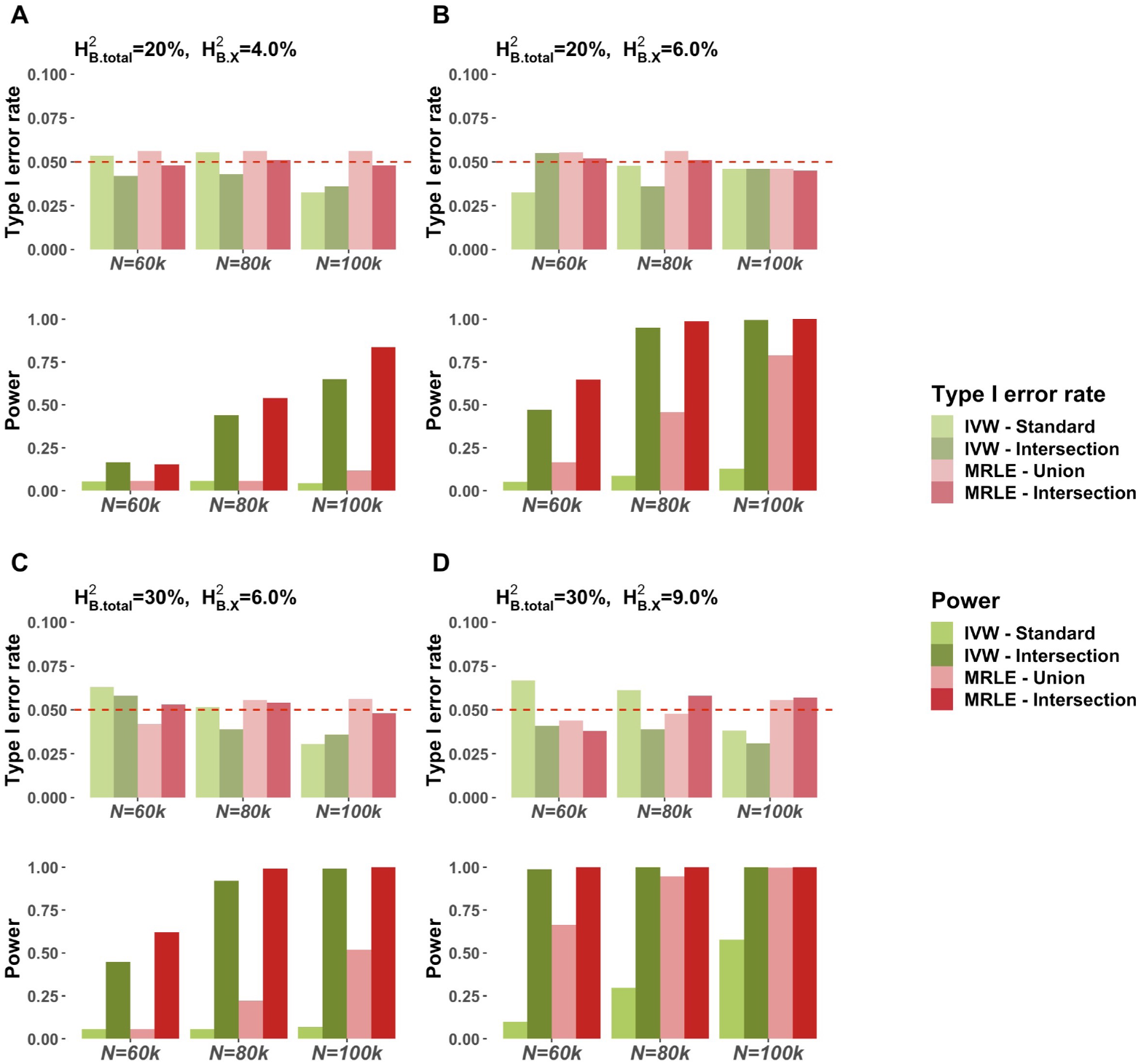
Simulation results assuming a total of *K* = 6 biomarkers based on 1000 simulations per setting. 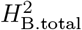 and 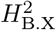 denote the total heritability of each biomarker and the heritability of each biomarker explained by the latent exposure, respectively. IVs are defined as either the SNPs associated with at least one biomarker (“IVW-Standard” and “MRLE-Union”, *α* = 5 × 10^-8^) or the SNPs associated with at least two biomarkers (“IVW-Intersection” and “MRLE-Intersection”, *α* = 5 × 10^-6^). In each subfigure, the upper panel shows the empirical type I error rate under *θ* = 0, and the lower panel shows the empirical power under *θ* = 0.1.

### 3.3 Simulations assuming correlated pleiotropy between biomarkers and the outcome

We next examined the performance of the tests under different types of pleiotropic effects. We first considered pleiotropy between the biomarkers and the outcome, where there exist SNPs that have correlated associations with the outcome and at least one of the biomarkers (Figure S1(A)). In our simulation, this was reflected by randomly assigning half of the *π_B_M* SNPs that had a direct effect on each biomarker to have another direct effect, *u_k,j_*, on the outcome, with mean 0, variance 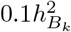, and *cor*(*γ_B_k,j__, u_k,j_*) = 0.15, *K* ∈ 1,…, *K*. We set *π_B_* = 2%, and for each biomarker, we set the total heritability to 0.3 or 0.4, and the proportion of heritability explained by the latent exposure to 0.15 or 0.2, which leads to a heritability of the biomarker explained by the latent exposure between 0.045 and 0.080.

In the presence of horizontal pleiotropy between a biomarker and the outcome, selecting SNPs significantly associated (i.e., *α* = 5 × 10^-8^) with any biomarker as IVs can lead to severe inflation in type I error rate, especially for the IVW test (Figure 4, “IVW-Standard”). Compared to the results in the no-pleiotropy scenario in Figures 3 and S4(B), at the same level of power, the probability of correctly identifying the causal direction also decreases for both tests with either of the two IV selection strategies (Figure S5(A)), although MRLE test still outperforms IVW test in terms of type I error control and identification of the direction of the effect. On the contrary, selecting SNPs that are associated with at least two biomarkers at a more liberal threshold (*α* = 5 × 10^-6^) as IVs can substantially improve the type I error control, power, and the probability of correcting identifying the direction of the effect for both methods. Under this strategy, MRLE has a consistently lower type I error rate that is close to the target level *α*_0_ = 0.05 and higher power compared to the IVW test.

**Fig. 4:**
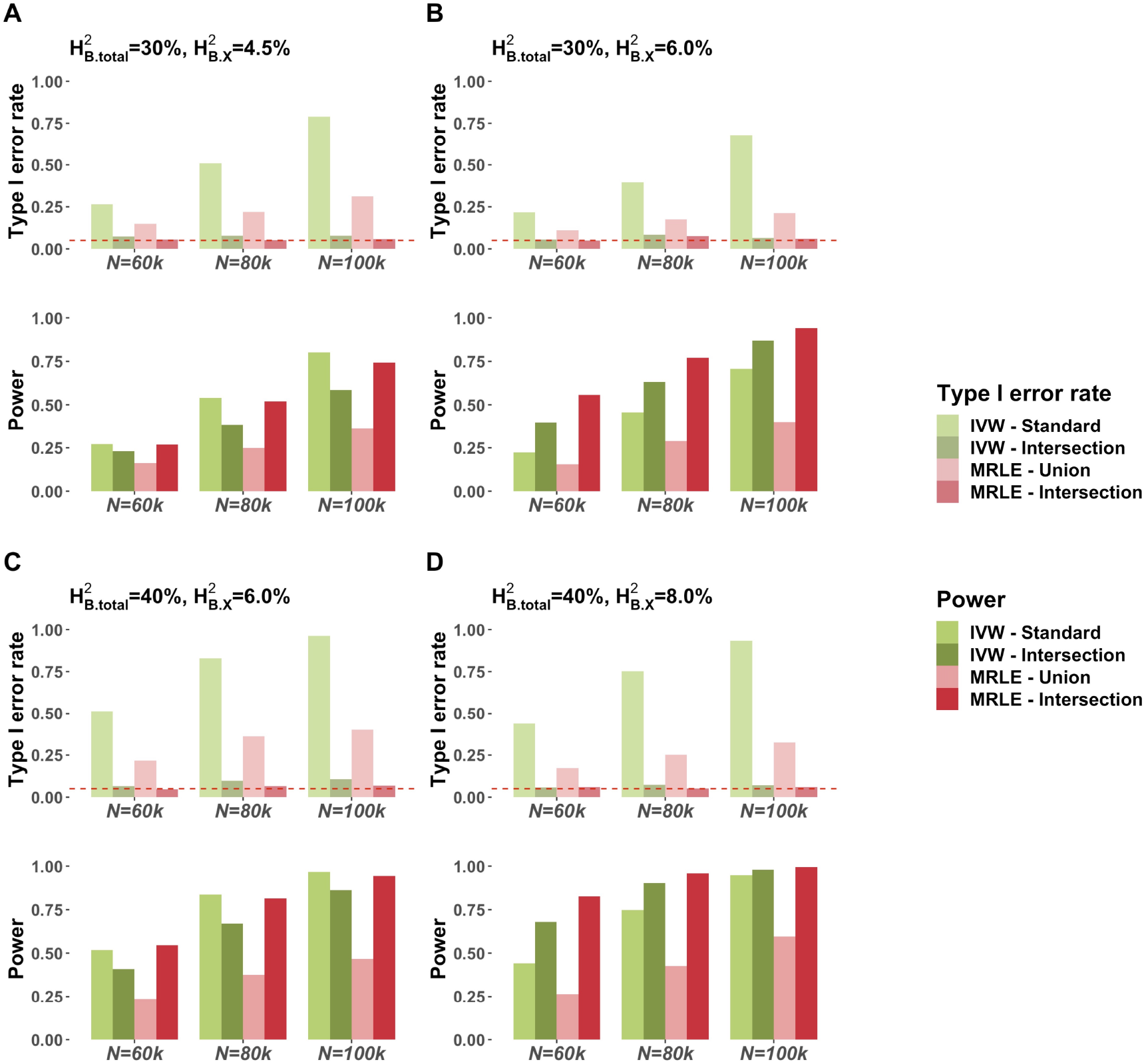
Simulation results assuming a total of *K* = 6 biomarkers and that there are SNPs that have correlated pleiotropic effects between some biomarkers and the outcome. 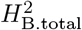 and 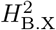 denote the total heritability of each biomarker and the heritability of each biomarker explained by the latent exposure, respectively. IVs are defined as either the SNPs associated with at least one biomarker (“IVW-Standard” and “MRLE-Union”, *α* = 5 × 10^-8^) or the SNPs associated with at least two biomarkers (“IVW-Intersection” and “MRLE-Intersection”, *α* = 5 × 10^-6^). In each subfigure, the upper panel shows the empirical type I error rate under *θ* = 0, and the lower panel shows the empirical power under *θ* = 0.1.

### 3.4 Simulations assuming correlated pleiotropy across biomarkers

Another type of pleiotropy that is likely to exist is the pleiotropy among biomarkers, where some SNPs have correlated direct effects across multiple biomarkers, which can be due to shared genetic pathways across biomarkers (Figure S1(B)). In the simulation, we introduced this type of pleiotropy by allowing 1/*K* of the *π_B_ M* SNPs that had a direct effect on biomarker *k (γ_k,j_*) to also have another direct effect on each of the other *K* – 1 biomarkers (*γ_l,j_, j* ≠ *k*), with mean 0, variances 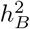, and cor(*γ_k,j_, γ_l,j_*) = 0.5, *k,l* ∈ 1,…, *K*. This leads to a total of *π_B_ M/K* SNPs to be directly associated with each individual biomarker, and *π_B_M/K* SNPs to be directly associated with each pair of biomarkers. We set *π_B_* = *K*%, and for each biomarker, we set the total heritability to 0.3 or 0.4, and the proportion of heritability explained by the latent exposure to 0.15 or 0.2, which leads to a heritability of the biomarker explained by the latent exposure between 0.045 and 0.08.

Simulation results show that in the presence of pleiotropy across biomarkers, strict type I error control can be achieved by either selecting SNPs associated with any biomarker (“IVW-Standard” and “MRLE-Union” in Figure 5) or selecting SNPs associated with at least two biomarkers (“IVW-Intersection” and “MRLE-Intersection” in Figure 5). Overall, the more stringent IV selection strategy (“IVW-Intersection” and “MRLE-Intersection”) yields a higher power and a higher probability of correctly identifying the causal direction for both IVW and MRLE tests (Figures 5 and S5(B)). Compared to IVW, MRLE shows a higher power and a higher probability of correctly identifying the causal direction using either of the two IV selection criteria.

**Fig. 5:**
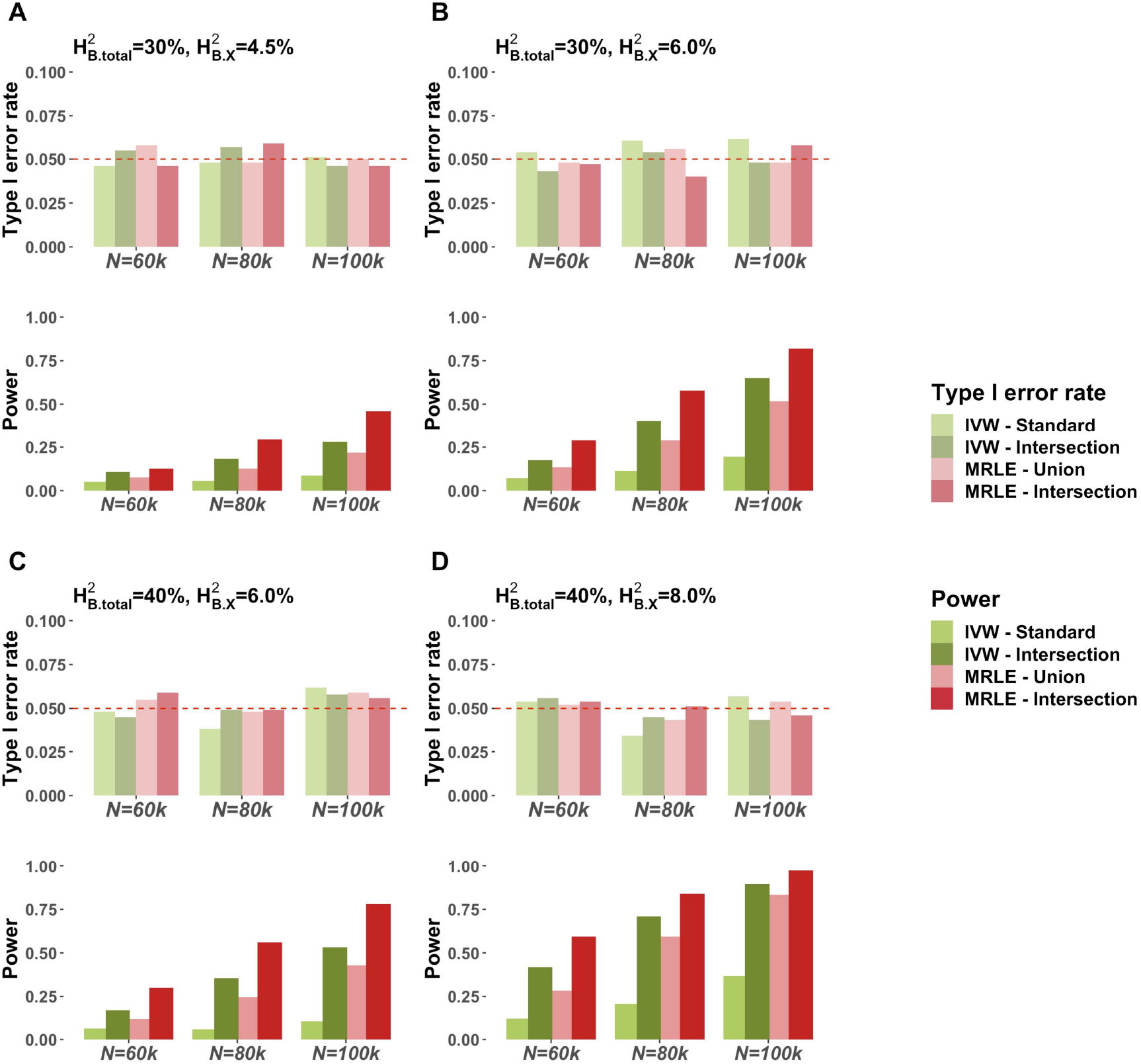
Simulation results assuming a total of *K* = 6 biomarkers and correlated pleiotropic effects across biomarkers. 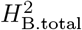 and 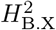 denote the total heritability of each biomarker and the heritability of each biomarker explained by the latent exposure, respectively. IVs are defined as either the SNPs associated with at least one biomarker (“IVW-Standard” and “MRLE-Union”, *α* = 5 × 10^-8^) or the SNPs associated with at least two biomarkers (“IVW-Intersection” and “MRLE-Intersection”, *α* = 5 × 10^-6^). In each subfigure, the upper panel shows the empirical type I error rate under *θ* = 0, and the lower panel shows the empirical power under *θ* = 0.1.

### 3.5 Simulations assuming correlated pleiotropy between the latent exposure and its biomarkers

In Section 3.4 we assess the validity and robustness of MRLE in the presence of correlated pleiotropy across biomarkers, i.e., there exist SNPs that have correlated direct effects, *γ*_1,*j*_,…, *γ_K,j_*, across the *K* biomarkers. Now we further consider a more complex scenario where in addition to the correlation structure among *γ_k,j_*s, there is also correlation between *γ_k,j_*s and *β_x,j_* the direct effect of SNP j on the latent exposure. In other words, there are SNPs that have pleiotropic effects described in Figure S1(B), or Figure S1(C), or both. Specifically, we conducted an additional simulation under the same simulation setting as in Section 3.2 except that we now only consider *K* = 6 biomarkers, and for each SNP *j* that had more than two nonzero effects among *β_x,j_, γ*_1,*j*_,…, *γ_K,j_*, we added an additional correlation 0.3 between any pair of the effects. Given that the number of SNPs with correlated direct effects on both the exposure and the biomarkers is small compared to the total number of causal SNPs, the heritability of the exposure and the biomarkers are approximately the same as that in the simulations in Section 3.2. Results in Figure 6 show that both IVW and MRLE have good type I error control under correlated pleiotropy across latent exposure and biomarkers. Both IVW and MRLE have higher power (Figure 6) and a higher probability of correctly identifying the causal direction (Figure S6) compared to the corresponding tests when there is no correlated pleiotropy across latent exposure and biomarkers (Figure 3, Figure S4(B)).

**Fig. 6:**
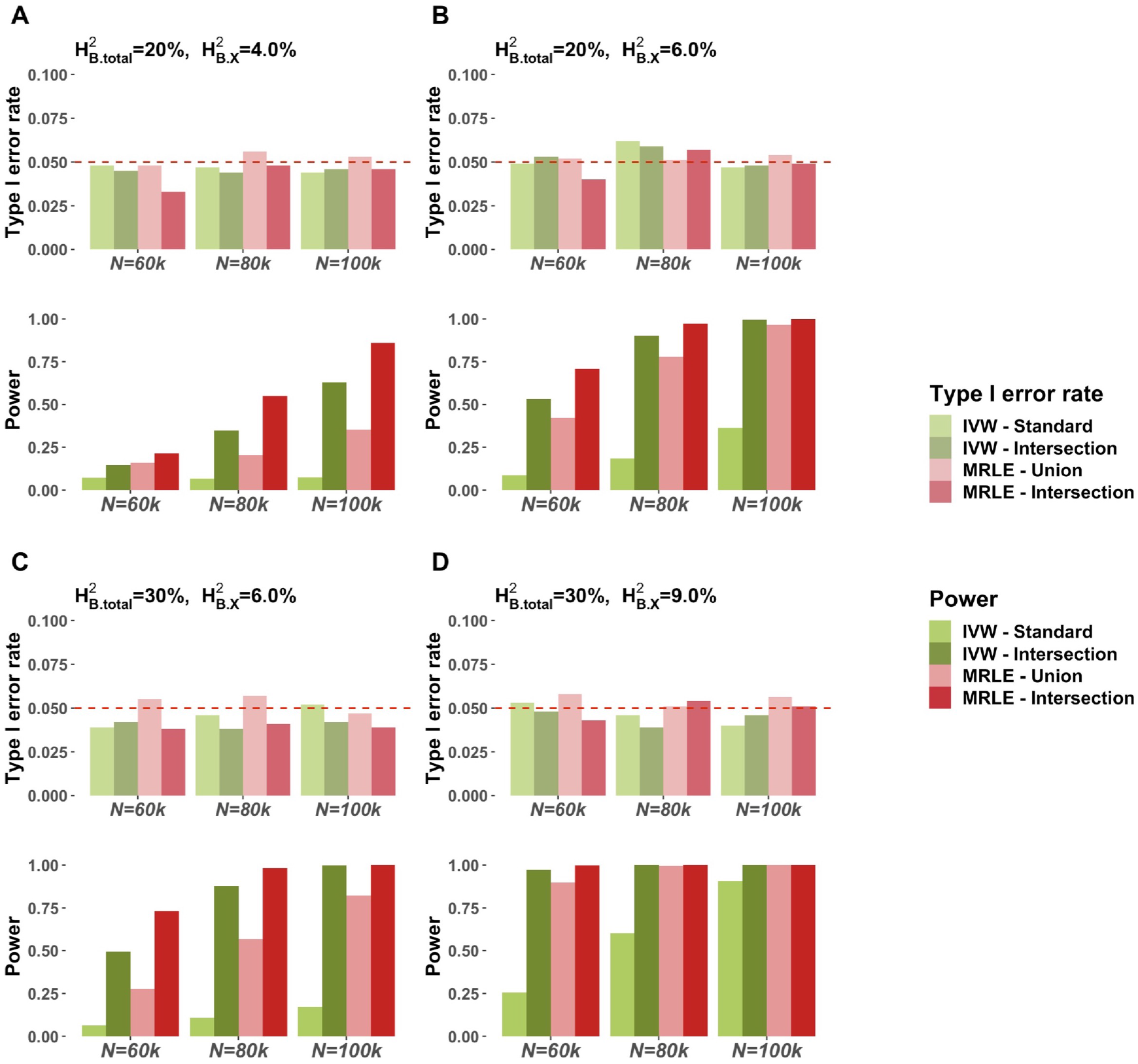
Simulation results assuming a total of *K* = 6 biomarkers and correlated pleiotropic effects across latent exposure and biomarkers. 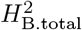 and 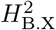 denote the total heritability of each biomarker and the heritability of each biomarker explained by the latent exposure, respectively. IVs are defined as either the SNPs associated with at least one biomarker (“IVW-Standard” and “MRLE-Union”, *α* = 5 × 10^-8^) or the SNPs associated with at least two biomarkers (“IVW-Intersection” and “MRLE-Intersection”, *α* = 5 × 10^-6^). In each subfigure, the upper panel shows the empirical type I error rate under *θ* = 0, and the lower panel shows the empirical power under *θ* = 0.1.

## 4. MR Analysis of Multiple Inflammatory Biomarkers on Risk of Five Diseases

Chronic inflammation has been long hypothesized to be one of the underlying causes of a spectrum of common diseases (Libby, 2007). Epidemiologic studies have used a variety of inflammation biomarkers to study the potential relationship between inflammation and disease risks (Hunter, 2012; Brenner *and others*, 2014; Bennett *and others*, 2018; Furman *and others*, 2019; Demir, 2020). In particular, C-reactive protein (CRP), a type of protein in blood produced by liver in response to inflammation, has been commonly used to associate chronic inflammation to a variety of diseases including heart disease (Collaboration *and others*, 2010; Shrivastava *and others*, 2015), ischemic stroke (Di Napoli *and others*, 2001; VanGilder *and others*, 2014), cancers of colon (Erlinger *and others*, 2004; Aleksandrova *and others*, 2010) and lung (Chaturvedi *and others*, 2010; Pastorino *and others*, 2017). However, recent MR studies have indicated that CRP itself is unlikely to be an underlying causal risk factor for these diseases. In addition to CRP, a variety of other biomarkers, including interleukin 6 (IL-6), interleukin 8 (IL-8), tumor necrosis factor alpha (TNF-α), and monocyte chemoattractant protein-1 (MCP-1), are commonly used to assess inflammation and hence associate with risks of diseases. MR analyses for these additional biomarkers, however, have been largely limited as sample sizes for the underlying GWAS have been typically fairly modest (Ahola-Olli *and others*, 2017; Höglund *and others*, 2019; Hillary *and others*, 2020; Russell *and others*, 2020).

Here we investigate the causal effect of chronic inflammation on a number of diseases including rheumatoid arthritis (RA), coronary artery disease (CAD), colorectal cancer (CRC), prostate cancer (PCa) and endometrial cancers (EC), all of which have been associated with one or more inflammatory biomarkers in previous studies (Choy and Panayi, 2001; Kraus and Arber, 2009; Friedenreich *and others*, 2013; Abu-Remaileh *and others*, 2015; Shrivastava *and others*, 2015; Izano *and others*, 2016; Platz *and others*, 2017; Li *and others*, 2018; Ridker *and others*, 2018; Cai *and others*, 2019; Subirana *and others*, 2018; Wang *and others*, 2019).

We apply the proposed MRLE method to test the causal effect of chronic inflammation on the diseases using summary-level data from publicly available GWAS for five commonly used systematic biomarkers of chronic inflammation, including, CRP, IL-6, IL-8, TNF-α and MCP-1. We ourselves generated the summary-level data for CRP by conducting a GWAS on 320041 unrelated, European-ancestry individuals in the UK Biobank who have CRP measurements available. We performed a GWAS across 1186957 common SNPs (minor allele frequency >5%) that are available in HapMap 3 (International HapMap 3 Consortium and others, 2010) based on additive genetic model adjusting for age, sex and body-mass index (BMI) using PLINK 2 (Chang *and others*, 2015; Purcell, S. M. and Chang, C. C., 2018). We used summary-level data for IL-6, IL-8, TNF-α and MCP-1 which were previously generated based on GWAS on up to 3596 European-ancestry participants in the Cardiovascular Risk in Young Finns Study (YFS) and up to 6313 European-ancestry participants in the FINRISK study that have the corresponding measurements available, after adjusting for age, sex, BMI and the first 10 genetic principal components (Ahola-Olli *and others*, 2017). Summary-level data for RA (Okada *and others*, 2014), CAD (Schunkert *and others*, 2011), CRC (Zhou *and others*, 2018), PCa (Schumacher *and others*, 2018), and EC (O’Mara *and others*, 2018) were all obtained from publicly available GWAS. As GWAS for CRC and EC have overlapping individuals with the UK Biobank-based GWAS for CRP, we excluded CRP from the set of biomarkers used in the analyses of CRC and EC. Detailed information on GWAS for the inflammatory biomarkers and diseases are summarized in Table S1.

To select IVs, we first conducted a filtering procedure (Zheng *and others*, 2017b) by removing the SNPs that were strand-ambiguous, had alleles that did not match those in the 1000 Genomes Project, or were within the major histocompatibility complex (MHC) region (26Mb - 34Mb on chromosome 6) since they may have complex large pleiotropic effects across multiple inflammation related traits (Trowsdale and Knight, 2013; Matzaraki *and others*, 2017). We then selected SNPs that were significantly associated with at least two of the inflammatory biomarkers. Considering the GWAS sample sizes, we used a more liberal instrument selection threshold, *α* = 10^-3^, for the four cytokine-type biomarkers (*N*_GWAS_ = 3454 ~ 8293), and a more stringent threshold, *α* = 5 × 10^-6^, for CRP (*N*_GWAS_ = 320041). To select independent SNPs, we conducted linkage disequilibrium (LD) clumping on the remaining SNPs with a window size *d* = 1MB and a cutoff for squared-correlation, *r*^2^ = 0.05 using PLINK (Purcell *and others*, 2007). Additionally, we removed SNPs that were significantly associated (α = 5 × 10^-8^) with the potential confounders of the association between inflammation and the outcomes (Rothenbacher *and others*, 2003), including systolic blood pressure (SBP), history of diabetes, smoking status, alcohol consumption status, high-density lipoprotein (HDL), and low-density lipoprotein (LDL), based on a GWAS we conducted on 320041 relatively unrelated European-ancestry UK Biobank individuals using PLINK 2 (Chang *and others*, 2015; Purcell, S. M. and Chang, C. C., 2018). These steps lead to a total of 53-58 IVs selected for the MR analyses on CRC and EC, and a total of 58-67 IVs for the MR analyses on CAD, RA and PCa.

Since summary-level data for the four cytokine-type biomarkers, IL-6, IL-8, TNF-*α* and MCP-1, were obtained from the same GWAS, we estimated the between-biomarker covariance (*c_k,k′_*s in the proposed estimating equations) by fitting bivariate LD score regressions as described in the Methods section. We also conducted fixed-effect IVW test on each biomarker, where the IVs were defined as the SNPs that were associated with the biomarker and at least one other biomarker. The MR analyses were conducted at a significance level of *α*_0_ = 0.05. The desired level of type I error rate for the single-biomarker IVW tests were set at 1 – (1 – *α*_0_)^1/*K*^, i.e., 0.0102 for CAD, RA and PCa, and 0.0127 for CRC and EC, to control FWER of the IVW test at 0.05 for each disease. Details of the IV selection procedure are summarized in Figure S7.

We observe that results on RA are significant based on all tests (Table 1). The proposed MRLE test detects a significant, positive effect of chronic inflammation (p-value=2.5 × 10^-27^). All single-biomarker IVW tests detect significant evidence as well, but the identified causal directions do not agree with each other: the tests based on IL-8, TNF-*α* and CRP suggest a positive effect of chronic inflammation on the risk of RA, while IL-6 and MCP-1 suggest a negative effect, thus no universal conclusion can be drawn for the effect on RA based on IVW tests. No single-biomarker IVW test shows significant evidence for a causal effect of chronic inflammation on the risk of CAD. Among the single-biomarker IVW tests, three of the biomarkers (CRP, TNF-α and IL-8) which indicate most significant evidence all seem to be associated with an increased risk of CAD. Similarly, no single-biomarker IVW test on CRC suggests a significant effect of chronic inflammation, although two of them (IL-8 and TNF-α) achieve borderline significance both indicating an increased risk of CRC being associated with higher level of inflammation. The proposed MRLE method, on the other hand, indicates a significant, positive effect of chronic inflammation on the risk of both CAD (p-value=0.012) and CRC (p-value=0.011). Neither the IVW tests nor the MRLE test detects any significant effect of chronic inflammation on PCa or EC.

**Table 1:**
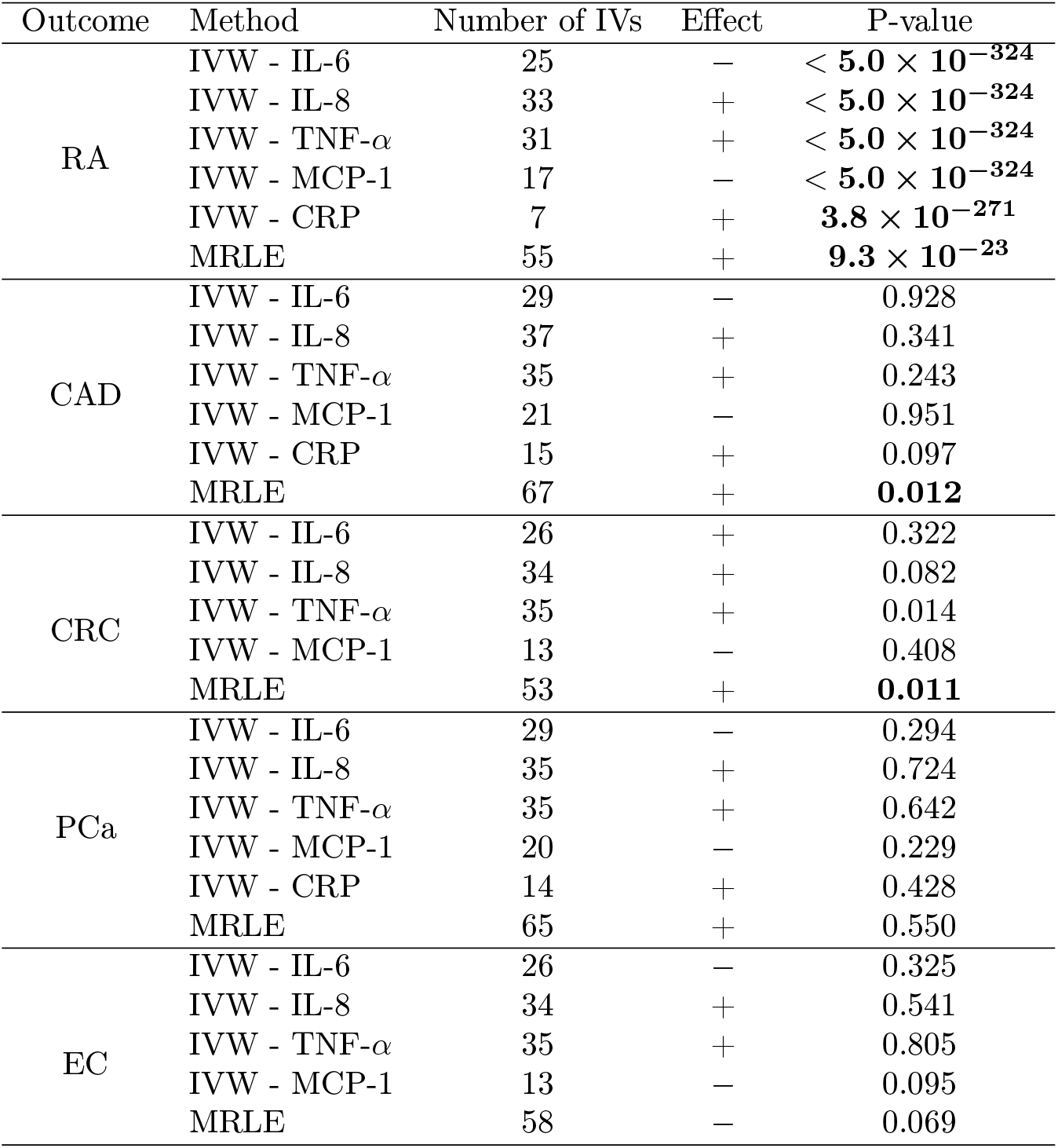
Causal effect of chronic inflammation on the risk of various diseases. The IVs used in MRLE are the SNPs associated with at least two of the inflammatory biomarkers. The IVs used in each single-biomarker IVW test are the SNPs associated with the biomarker and at least one other biomarker. Significance threshold for IV selection is set to *α* = 5 × 10^-6^ for CRP and *α* = 10^-3^ for the other biomarkers. Bold font indicates significant conclusion. Significance thresholds for IVW tests are set to 1 – (1 – *α*_0_)^1/*K*^, i.e., 0.0102 for CAD, RA and PCa, and 0.0127 for CRC and EC, to control FWER at 0.05. CRP were excluded from the tests on CRC and EC due to overlapping individuals in GWAS.

| Outcome | Method | Number of IVs | Effect | P-value |
| --- | --- | --- | --- | --- |
| RA | IVW - IL-6 | 25 | — | $< \mathbf{5.0 \times 10^{-324}}$ |
| | IVW - IL-8 | 33 | + | $< \mathbf{5.0 \times 10^{-324}}$ |
| | IVW - TNF-α | 31 | + | $< \mathbf{5.0 \times 10^{-324}}$ |
| | IVW - MCP-1 | 17 | — | $< \mathbf{5.0 \times 10^{-324}}$ |
| | IVW - CRP | 7 | + | $\mathbf{3.8 \times 10^{-271}}$ |
| | MRLE | 55 | + | $\mathbf{9.3 \times 10^{-23}}$ |
| CAD | IVW - IL-6 | 29 | — | 0.928 |
|  | IVW - IL-8 | 37 | + | 0.341 |
| | IVW - TNF-α | 35 | + | 0.243 |
|  | IVW - MCP-1 | 21 | — | 0.951 |
|  | IVW - CRP | 15 | + | 0.097 |
|  | MRLE | 67 | + | **0.012** |
| CRC | IVW - IL-6 | 26 | + | 0.322 |
|  | IVW - IL-8 | 34 | + | 0.082 |
| | IVW - TNF-α | 35 | + | 0.014 |
|  | IVW - MCP-1 | 13 | — | 0.408 |
|  | MRLE | 53 | + | **0.011** |
| PCa | IVW - IL-6 | 29 | — | 0.294 |
|  | IVW - IL-8 | 35 | + | 0.724 |
| | IVW - TNF-α | 35 | + | 0.642 |
|  | IVW - MCP-1 | 20 | — | 0.229 |
|  | IVW - CRP | 14 | + | 0.428 |
|  | MRLE | 65 | + | 0.550 |
| EC | IVW - IL-6 | 26 | — | 0.325 |
|  | IVW - IL-8 | 34 | + | 0.541 |
| | IVW - TNF-α | 35 | + | 0.805 |
|  | IVW - MCP-1 | 13 | — | 0.095 |
|  | MRLE | 58 | — | 0.069 |

## 5. Discussion

We propose a novel method for MR analysis for testing the causal effect of an unobservable latent exposure utilizing multiple traits co-regulated by the exposure. Through a set of extensive simulation studies and data analyses, we demonstrate that the proposed method overcomes various challenges associated with the standard MR analyses that use individual observable traits associated with the latent exposure and their associated genetic instruments.

Several practical issues merit consideration. First, the validity of the selected observable traits as surrogates for the latent exposure of interest. Theoretically, including more observable traits for latent exposure can provide higher power. However, the inclusion of invalid traits, i.e., traits that are actually not regulated by the latent exposure or/and themselves have a direct causal effect on the exposure, can affect both the type I error rate and power of the tests. Second, a strict IV selection procedure is crucial. Other than the commonly implemented filtering procedures such as removing SNPs that are significantly associated with the potential confounders, we recommend selecting the SNPs that are associated with at least two traits. We have shown by simulation that this more strict criterion can efficiently reduce the number of invalid IVs selected, thus providing higher power and more strict type I error control under various types of pleiotropy. Additionally, although a more liberal significance threshold may be used for an individual trait when the sample size is relatively small, there needs to be rigorous criteria to ensure the selection of valid instruments. Recent methods for discovering strictly pleiotropic associations across multiple traits can be potentially used for instrument selection (Ray and Chatterjee, 2020).

Our study has several limitations. First, our current model assumes no horizontal pleiotropy between the latent exposure and the outcome. There has been a considerable amount of research in the recent past on weakening the no horizontal pleiotropy assumption when the exposure is directly observable (Bowden *and others*, 2016; Hartwig *and others*, 2017; Verbanck *and others*, 2018; Qi and Chatterjee, 2019; Burgess *and others*, 2020). Our method, on the other hand, provides the formal framework for carrying an MR analysis for a latent exposure based on multiple biomarkers. The method is robust to pleiotropic effects of genetic variants across biomarkers, between the latent exposure and the biomarkers, and between the biomarkers and the outcome. Future studies are merited to explore how MRLE can be further strengthened to take into account possible horizontal pleiotropy between the latent exposure and the outcome.

Results from our MR analysis of chronic inflammation and various diseases should be interpreted cautiously. First, null results for certain diseases, such as PCa and EC, may be due to the inability of the set of the biomarkers used to capture relevant aspects of inflammation. Second, the GWAS sample for all the inflammatory biomarkers except CRP was relatively small, causing substantially large uncertainty of the effect estimates. In the future, our results need to be confirmed using results from much larger GWAS of inflammatory biomarkers when such data become available.

## Supporting information

Supplementary Information including Web Appendices and Supplementary Figures.

## 6. Software

The R package “MRLE” and the R code for simulations and data analyses are available at https://github.com/Jin93/MRLE.

## 7. Supplementary Materials

The reader is referred to the on-line Supplementary Materials for technical appendices and additional simulation results.

## 8. Acknowledgements

The UK Biobank data was accessed via application ID 17712. The research of Drs. Jin Jin and Nilanjan Chatterjee were supported by an R01 grant from the National Human Genome Research Institute R01HG010480. The research of Dr. Jin Jin was also supported by a K99 grant from the National Human Genome Research Institute K99HG012223.

## References

Abifadel, Marianne, Varret, Mathilde, Rabès, Jean-Pierre, Allard, Delphine, Ouguerram, Khadija, Devillers, Martine, Cruaud, Corinne, Benjannet, Suzanne, Wickham, Louise, Erlich, Danièle and others. (2003). Mutations in pcsk9 cause autosomal dominant hypercholesterolemia. Nature genetics 34(2), 154–156.

Abu-Remaileh, Monther, Bender, Sebastian, Raddatz, Günter, Ansari, Ihab, Cohen, Daphne, Gutekunst, Julian, Musch, Tanja, Linhart, Heinz, Breiling, Achim, Pikarsky, Eli and others. (2015). Chronic inflammation induces a novel epigenetic program that is conserved in intestinal adenomas and in colorectal cancer. Cancer research 75(10), 2120–2130.

Ahola-Olli, Ari V, Würtz, Peter, Havulinna, Aki S, Aalto, Kristiina, Pitkänen, Niina, Lehtimäki, Terho, Kähönen, Mika, Lyytikäinen, Leo-Pekka, Raitoharju, Emma, Seppälä, Ilkka and others. (2017). Genome-wide association study identifies 27 loci influencing concentrations of circulating cytokines and growth factors. The American Journal of Human Genetics 100(1), 40–50.

Aleksandrova, Krasimira, Jenab, Mazda, Boeing, Heiner, Jansen, Eugene, Bueno-de Mesquita, H Bas, Rinaldi, Sabina, Riboli, Elio, Overvad, Kim, Dahm, Christina C, Olsen, Anja and others. (2010). Circulating c-reactive protein concentrations and risks of colon and rectal cancer: a nested case-control study within the european prospective investigation into cancer and nutrition. American journal of epidemiology 172(4), 407–418.

Bautista, Leonelo E, Smeeth, Liam, Hingorani, Aroon D and Casas, Juan P. (2006). Estimation of bias in nongenetic observational studies using “mendelian triangulation”. Annals of epidemiology 16(9), 675–680.

Bennett, Jeanette M, Reeves, Glenn, Billman, George E and Sturmberg, Joachim P. (2018). Inflammation–nature’s way to efficiently respond to all types of challenges: implications for understanding and managing “the epidemic” of chronic diseases. Frontiers in Medicine 5, 316.

Black, Louise, Panayiotou, Margarita and Humphrey, Neil. (2019). The dimensionality and latent structure of mental health difficulties and wellbeing in early adolescence. PloS one 14(2), e0213018.

Bowden, Jack, Davey Smith, George and Burgess, Stephen. (2015). Mendelian randomization with invalid instruments: effect estimation and bias detection through egger regression. International journal of epidemiology 44(2), 512–525.

Bowden, Jack, Davey Smith, George, Haycock, Philip C and Burgess, Stephen. (2016). Consistent estimation in mendelian randomization with some invalid instruments using a weighted median estimator. Genetic epidemiology 40(4), 304–314.

Brenner, Darren R, Scherer, Dominique, Muir, Kenneth, Schildkraut, Joellen, Boffetta, Paolo, Spitz, Margaret R, Le Marchand, Loic, Chan, Andrew T, Goode, Ellen L, Ulrich, Cornelia M and others. (2014). A review of the application of inflammatory biomarkers in epidemiologic cancer research. Cancer Epidemiology and Prevention Biomarkers 23(9), 1729–1751.

Burgess, Stephen, Foley, Christopher N, Allara, Elias, Staley, James R and Howson, Joanna MM. (2020). A robust and efficient method for mendelian randomization with hundreds of genetic variants. Nature communications 11(1), 376.

Burgess, Stephen and Thompson, Simon G. (2015). Multivariable mendelian randomization: the use of pleiotropic genetic variants to estimate causal effects. American journal of epidemiology 181(4), 251–260.

Cai, Tommaso, Santi, Raffaella, Tamanini, Irene, Galli, Ilaria Camilla, Perletti, Gianpaolo, Bjerklund Johansen, Truls E and Nesi, Gabriella. (2019). Current knowledge of the potential links between inflammation and prostate cancer. International journal of molecular sciences 20(15), 3833.

Chang, Christopher C, Chow, Carson C, Tellier, Laurent CAM, Vattikuti, Shashaank, Purcell, Shaun M and Lee, James J. (2015). Second-generation plink: rising to the challenge of larger and richer datasets. Gigascience 4(1), s13742–015.

Chaturvedi, Anil K, Caporaso, Neil E, Katki, Hormuzd A, Wong, Hui-Lee, Chatterjee, Nilanjan, Pine, Sharon R, Chanock, Stephen J, Goedert, James J and Engels, Eric A. (2010). C-reactive protein and risk of lung cancer. Journal of clinical oncology 28(16), 2719.

Choy, Ernest HS and Panayi, Gabriel S. (2001). Cytokine pathways and joint inflammation in rheumatoid arthritis. New England Journal of Medicine 344(12), 907–916.

Cinelli, Carlos, LaPierre, Nathan, Hill, Brian L, Sankararaman, Sriram and Eskin, Eleazar. (2022). Robust mendelian randomization in the presence of residual population stratification, batch effects and horizontal pleiotropy. Nature communications 13(1), 1–13.

Cohen, Jonathan C, Boerwinkle, Eric, Mosley Jr, Thomas H and Hobbs, Helen H. (2006). Sequence variations in pcsk9, low ldl, and protection against coronary heart disease. New England Journal of Medicine 354(12), 1264–1272.

Collaboration, Emerging Risk Factors and others. (2010). C-reactive protein concentration and risk of coronary heart disease, stroke, and mortality: an individual participant meta-analysis. The Lancet 375(9709), 132–140.

Davey Smith, George and Ebrahim, Shah. (2003). ‘mendelian randomization’: can genetic epidemiology contribute to understanding environmental determinants of disease? International journal of epidemiology 32(1), 1–22.

Demir, Sedat. (2020). The process of acute and chronic inflammation: Biomarkers and their relationship with diseases. In: Role of Nutrition in Providing Pro-/Anti-Inflammatory Balance: Emerging Research and Opportunities. IGI Global, pp. 1–23.

Di Napoli, Mario, Papa, Francesca and Bocola, Vittorio. (2001). C-reactive protein in ischemic stroke: an independent prognostic factor. Stroke 32(4), 917–924.

Emdin, Connor A, Khera, Amit V and Kathiresan, Sekar. (2017). Mendelian randomization. Jama 318(19), 1925–1926.

Erlinger, Thomas P, Platz, Elizabeth A, Rifai, Nader and Helzlsouer, Kathy J. (2004). C-reactive protein and the risk of incident colorectal cancer. Jama 291(5), 585–590.

Evans, David M and Davey Smith, George. (2015). Mendelian randomization: new applications in the coming age of hypothesis-free causality. Annual review of genomics and human genetics 16, 327–350.

Friedenreich, Christine M, Langley, Annie R, Speidel, Thomas P, Lau, David CW, Courneya, Kerry S, Csizmadi, Ilona, Magliocco, Anthony M, Yasui, Yutaka and Cook, Linda S. (2013). Case–control study of inflammatory markers and the risk of endometrial cancer. European journal of cancer prevention 22(4), 374–379.

Furman, David, Campisi, Judith, Verdin, Eric, Carrera-Bastos, Pedro, Targ, Sasha, Franceschi, Claudio, Ferrucci, Luigi, Gilroy, Derek W, Fasano, Alessio, Miller, Gary W and others. (2019). Chronic inflammation in the etiology of disease across the life span. Nature medicine 25(12), 1822–1832.

Greenland, Sander. (2000). An introduction to instrumental variables for epidemiologists. International journal of epidemiology 29(4), 722–729.

Gu, Qianqian, Spinelli, John J, Dummer, Trevor BJ, McDonald, Treena E, Moore, Steven C and Murphy, Rachel A. (2018). Metabolic profiling of adherence to diet, physical activity and body size recommendations for cancer prevention. Scientific reports 8(1), 1–11.

Hall, Alastair R and others. (2005). Generalized method of moments. Oxford university press.

Hansen, Lars Peter. (1982). Large sample properties of generalized method of moments estimators. Econometrica: Journal of the Econometric Society, 1029–1054.

Hartwig, Fernando Pires, Davey Smith, George and Bowden, Jack. (2017). Robust inference in summary data mendelian randomization via the zero modal pleiotropy assumption. International journal of epidemiology 46(6), 1985–1998.

Hillary, Robert F, Trejo-Banos, Daniel, Kousathanas, Athanasios, McCartney, Daniel L, Harris, Sarah E, Stevenson, Anna J, Patxot, Marion, Ojavee, Sven Erik, Zhang, Qian, Liewald, David C and others. (2020). Multi-method genome- and epigenome-wide studies of inflammatory protein levels in healthy older adults. Genome medicine 12(1), 1–15.

Höglund, Julia, Rafati, Nima, Rask-Andersen, Mathias, Enroth, Stefan, Karlsson, Torgny, Ek, Weronica E and Johansson, Åsa. (2019). Improved power and precision with whole genome sequencing data in genome-wide association studies of inflammatory biomarkers. Scientific reports 9(1), 1–14.

Hunter, Philip. (2012). The inflammation theory of disease: The growing realization that chronic inflammation is crucial in many diseases opens new avenues for treatment. EMBO reports 13(11), 968–970.

Izano, Monika, Wei, Esther K, Tai, Caroline, Swede, Helen, Gregorich, Steven, Harris, Tamara B, Klepin, Heidi, Satterfield, Suzanne, Murphy, Rachel, Newman, Anne B and others. (2016). Chronic inflammation and risk of colorectal and other obesity-related cancers: the health, aging and body composition study. International journal of cancer 138(5), 1118–1128.

Kraus, Sarah and Arber, Nadir. (2009). Inflammation and colorectal cancer. Current opinion in pharmacology 9(4), 405–410.

Lawlor, Debbie A, Harbord, Roger M, Sterne, Jonathan AC, Timpson, Nic and Davey Smith, George. (2008). Mendelian randomization: using genes as instruments for making causal inferences in epidemiology. Statistics in medicine 27(8), 1133–1163.

Levey, Andrew S, Stevens, Lesley A, Schmid, Christopher H, Zhang, Yaping, Castro III, Alejandro F, Feldman, Harold I, Kusek, John W, Eggers, Paul, Van Lente, Frederick, Greene, Tom and others. (2009). A new equation to estimate glomerular filtration rate. Annals of internal medicine 150(9), 604–612.

Li, Hongyu, Sun, Kai, Zhao, Ruiping, Hu, Jiang, Hao, Zhiru, Wang, Fei, Lu, Yaojun, Liu, Fu and Zhang, Yong. (2018). Inflammatory biomarkers of coronary heart disease. Front Biosci (Schol Ed) 10, 185–96.

Matzaraki, Vasiliki, Kumar, Vinod, Wijmenga, Cisca and Zhernakova, Alexandra. (2017). The mhc locus and genetic susceptibility to autoimmune and infectious diseases. Genome biology 18(1), 76.

Morrison, Jean, Knoblauch, Nicholas, Marcus, Joseph H, Stephens, Matthew and He, Xin. (2020). Mendelian randomization accounting for correlated and uncorrelated pleiotropic effects using genome-wide summary statistics. Nature genetics 52(7), 740–747.

Newey, Whitney K and McFadden, Daniel. (1994). Large sample estimation and hypothesis testing. Handbook of econometrics 4, 2111–2245.

Okada, Yukinori, Wu, Di, Trynka, Gosia, Raj, Towfique, Terao, Chikashi, Ikari, Katsunori, Kochi, Yuta, Ohmura, Koichiro, Suzuki, Akari, Yoshida, Shinji and others. (2014). Genetics of rheumatoid arthritis contributes to biology and drug discovery. Nature 506(7488), 376–381.

Oluwagbemigun, Kolade, Foerster, Jana, Watkins, Claire, Fouhy, Fiona, Stanton, Catherine, Bergmann, Manuela M, Boeing, Heiner and Nöthlings, Ute. (2020). Dietary patterns are associated with serum metabolite patterns and their association is influenced by gut bacteria among older german adults. The Journal of nutrition 150(1), 149–158.

O’Mara, Tracy A, Glubb, Dylan M, Amant, Frederic, Annibali, Daniela, Ashton, Katie, Attia, John, Auer, Paul L, Beckmann, Matthias W, Black, Amanda, Bolla, Manjeet K and others. (2018). Identification of nine new susceptibility loci for endometrial cancer. Nature communications 9(1), 1–12.

Pastorino, Ugo, Morelli, Daniele, Leuzzi, Giovanni, Gisabella, Mara, Suatoni, Paola, Taverna, Francesca, Bertocchi, Elena, Boeri, Mattia, Sozzi, Gabriella, Cantarutti, Anna and others. (2017). Baseline and postoperative c-reactive protein levels predict mortality in operable lung cancer. European journal of cancer 79, 90–97.

Pierce, Brandon L, Ahsan, Habibul and VanderWeele, Tyler J. (2011). Power and instrument strength requirements for mendelian randomization studies using multiple genetic variants. International journal of epidemiology 40(3), 740–752.

Pierce, Brandon L and Burgess, Stephen. (2013). Efficient design for mendelian randomization studies: subsample and 2-sample instrumental variable estimators. American journal of epidemiology 178(7), 1177–1184.

Platz, Elizabeth A, Kulac, Ibrahim, Barber, John R, Drake, Charles G, Joshu, Corinne E, Nelson, William G, Lucia, M Scott, Klein, Eric A, Lippman, Scott M, Parnes, Howard L and others. (2017). A prospective study of chronic inflammation in benign prostate tissue and risk of prostate cancer: linked pcpt and select cohorts. Cancer Epidemiology and Prevention Biomarkers 26(10), 1549–1557.

Purcell, Shaun, Neale, Benjamin, Todd-Brown, Kathe, Thomas, Lori, Ferreira, Manuel AR, Bender, David, Maller, Julian, Sklar, Pamela, De Bakker, Paul IW, Daly, Mark J and others. (2007). Plink: a tool set for whole-genome association and population-based linkage analyses. The American journal of human genetics 81(3), 559–575.

Purcell, S. M. and Chang, C. C. (2018). PLINK 2.0. https://www.cog-genomics.org/plink/2.0/. Accessed: 2020-05-15.

Qi, Guanghao and Chatterjee, Nilanjan. (2018). Heritability informed power optimization (hipo) leads to enhanced detection of genetic associations across multiple traits. PLoS genetics 14(10), e1007549.

Qi, Guanghao and Chatterjee, Nilanjan. (2019). Mendelian randomization analysis using mixture models for robust and efficient estimation of causal effects. Nature communications 10(1), 1–10.

Qi, Guanghao and Chatterjee, Nilanjan. (2021). A comprehensive evaluation of methods for mendelian randomization using realistic simulations and an analysis of 38 biomarkers for risk of type 2 diabetes. International journal of epidemiology 50(4), 1335–1349.

Qian, Shehua, Golubnitschaja, Olga and Zhan, Xianquan. (2019). Chronic inflammation: key player and biomarker-set to predict and prevent cancer development and progression based on individualized patient profiles. Epma Journal 10(4), 365–381.

Ridker, Paul M, MacFadyen, Jean G, Everett, Brendan M, Libby, Peter, Thuren, Tom, Glynn, Robert J, Kastelein, John, Koenig, Wolfgang, Genest, Jacques, Lorenzatti, Alberto and others. (2018). Relationship of c-reactive protein reduction to cardiovascular event reduction following treatment with canakinumab: a secondary analysis from the cantos randomised controlled trial. The Lancet 391(10118), 319–328.

Rothenbacher, Dietrich, Brenner, Hermann, Hoffmeister, Albrecht, Mertens, Thomas, Persson, Kenneth and Koenig, Wolfgang. (2003). Relationship between infectious burden, systemic inflammatory response, and risk of stable coronary artery disease: role of confounding and reference group. Atherosclerosis 170(2), 339–345.

Russell, Abigail Emma, Ford, Tamsin, Gunnell, David, Heron, Jon, Joinson, Carol, Moran, Paul, Relton, Caroline, Suderman, Matthew, Hemani, Gibran, and Mars, Becky. (2020). Investigating evidence for a causal association between inflammation and self-harm: a multivariable mendelian randomisation study. Brain, behavior, and immunity 89, 43–50.

Schumacher, Fredrick R, Al Olama, Ali Amin, Berndt, Sonja I, Benlloch, Sara, Ahmed, Mahbubl, Saunders, Edward J, Dadaev, Tokhir, Leongamornlert, Daniel, Anokian, Ezequiel, Cieza-Borrella, Clara and others. (2018). Association analyses of more than 140,000 men identify 63 new prostate cancer susceptibility loci. Nature genetics 50(7), 928–936.

Schunkert, Heribert, König, Inke R, Kathiresan, Sekar, Reilly, Muredach P, Assimes, Themistocles L, Holm, Hilma, Preuss, Michael, Stewart, Alexandre FR, Barbalic, Maja, Gieger, Christian and others. (2011). Large-scale association analysis identifies 13 new susceptibility loci for coronary artery disease. Nature genetics 43(4), 333–338.

Shapland, Chin Yang, Zhao, Qingyuan and Bowden, Jack. (2022). Profile-likelihood bayesian model averaging for two-sample summary data mendelian randomization in the presence of horizontal pleiotropy. Statistics in Medicine 41(6), 1100–1119.

Shrivastava, Amit Kumar, Singh, Harsh Vardhan, Raizada, Arun and Singh, Sanjeev Kumar. (2015). C-reactive protein, inflammation and coronary heart disease. The Egyptian Heart Journal 67(2), 89–97.

Sproston, Nicola R and Ashworth, Jason J. (2018). Role of c-reactive protein at sites of inflammation and infection. Frontiers in immunology 9, 754.

Subirana, Isaac, Fitó, Montserrat, Diaz, Oscar, Vila, Joan, Francés, Albert, Delpon, Eva, Sanchis, Juan, Elosua, Roberto, Muñoz-Aguayo, Daniel, Dégano, Irene R and others. (2018). Prediction of coronary disease incidence by biomarkers of inflammation, oxidation, and metabolism. Scientific reports 8(1), 1–7.

Trowsdale, John and Knight, Julian C. (2013). Major histocompatibility complex genomics and human disease. Annual review of genomics and human genetics 14, 301–323.

Tsanas, Athanasios, Saunders, Kate, Bilderbeck, Amy, Palmius, Niclas, Goodwin, Guy and De Vos, Maarten. (2017). Clinical insight into latent variables of psychiatric questionnaires for mood symptom self-assessment. JMIR mental health 4(2), e15.

Turley, Patrick, Walters, Raymond K, Maghzian, Omeed, Okbay, Aysu, Lee, James J, Fontana, Mark Alan, Nguyen-Viet, Tuan Anh, Wedow, Robbee, Zacher, Meghan, Furlotte, Nicholas A and others. (2018). Multi-trait analysis of genome-wide association summary statistics using mtag. Nature genetics 50(2), 229–237.

VanGilder, Reyna L, Davidov, Danielle M, Stinehart, Kyle R, Huber, Jason D, Turner, Ryan C, Wilson, Karen S, Haney, Eric, Davis, Stephen M, Chantler, Paul D, Theeke, Laurie and others. (2014). C-reactive protein and long-term ischemic stroke prognosis. Journal of Clinical Neuroscience 21(4), 547–553.

Verbanck, Marie, Chen, Chia-yen, Neale, Benjamin and Do, Ron. (2018). Detection of widespread horizontal pleiotropy in causal relationships inferred from mendelian randomization between complex traits and diseases. Nature genetics 50(5), 693–698.

Wang, Xiaoliang, Dai, James Y, Albanes, Demetrius, Arndt, Volker, Berndt, Sonja I, Bézieau, Stéphane, Brenner, Hermann, Buchanan, Daniel D, Butterbach, Katja, Caan, Bette and others. (2019). Mendelian randomization analysis of c-reactive protein on colorectal cancer risk. International journal of epidemiology 48(3), 767–780.

Yu, Zhi, Coresh, Josef, Qi, Guanghao, Grams, Morgan, Boerwinkle, Eric, Snieder, Harold, Teumer, Alexander, Pattaro, Cristian, Köttgen, Anna, Chatterjee, Nilanjan and others. (2020). A bidirectional mendelian randomization study supports causal effects of kidney function on blood pressure. Kidney international 98(3), 708–716.

Zhao, Qingyuan, Wang, Jingshu, Hemani, Gibran, Bowden, Jack, Small, Dylan S and others. (2020). Statistical inference in two-sample summary-data mendelian randomization using robust adjusted profile score. Annals of Statistics 48(3), 1742–1769.

Zheng, Jie, Baird, Denis, Borges, Maria-Carolina, Bowden, Jack, Hemani, Gibran, Haycock, Philip, Evans, David M and Smith, George Davey. (2017a). Recent developments in mendelian randomization studies. Current epidemiology reports 4(4), 330–345.

Zheng, Jie, Erzurumluoglu, A Mesut, Elsworth, Benjamin L, Kemp, John P, Howe, Laurence, Haycock, Philip C, Hemani, Gibran, Tansey, Katherine, Laurin, Charles, Pourcain, Beate St and others. (2017b). Ld hub: a centralized database and web interface to perform ld score regression that maximizes the potential of summary level gwas data for snp heritability and genetic correlation analysis. Bioinformatics 33(2), 272–279.

Zhou, Wei, Nielsen, Jonas B, Fritsche, Lars G, Dey, Rounak, Gabrielsen, Maiken E, Wolford, Brooke N, LeFaive, Jonathon, VandeHaar, Peter, Gagliano, Sarah A, Gifford, Aliya and others. (2018). Efficiently controlling for case-control imbalance and sample relatedness in large-scale genetic association studies. Nature genetics 50(9), 1335–1341.

Zhu, Zhihong, Zheng, Zhili, Zhang, Futao, Wu, Yang, Trzaskowski, Maciej, Maier, Robert, Robinson, Matthew R, McGrath, John J, Visscher, Peter M, Wray, Naomi R and others. (2018). Causal associations between risk factors and common diseases inferred from gwas summary data. Nature communications 9(1), 1–12.

