## Supplementary Information including Web Appendices and Supplementary Figures. for "Mendelian Randomization Analysis Using Multiple Biomarkers of an Underlying Common Exposure"

### APPENDIX A: ASYMPTOTIC BIAS OF THE STANDARD IVW ESTIMATOR

While there is currently no established method for testing the causal effect of a latent exposure, there have been attempts which test the causal effect of the biomarkers of the exposure as an alternative. A simple example is an approach which tests the effect of a biomarker of $X$ of $Y$ using the commonly implemented fixed-effect inverse-variance weighted (IVW) estimator.[1] Under the NO Measurement Error (NOME) assumption, i.e. $Var(\widehat{\beta}_{B_{k,j}}) = 0$, and the assumption that $\sigma^2_{Y,j}$s are fixed, the IVW estimator for $\theta$ using a single biomarker, $B$, is defined as

$$\widehat{\theta}^{IVW} = \frac{\sum_{j=1}^{M} \frac{\widehat{\beta}_{Y,j}\widehat{\beta}_{B,j}}{\sigma^2_{Y,j}}}{\sum_{j=1}^{M} \frac{\widehat{\beta}^2_{B,j}}{\sigma^2_{Y,j}}},$$

with a standard error $\left(\sum_{j=1}^{M}\left(\widehat{\beta}^2_{B,j}/\sigma^2_{Y,j}\right)\right)^{-1/2}$.

Assume that among the $M$ IVs selected for the biomarker, the first $p_v M$ are valid IVs for the latent exposure, while the rest are invalid which are associated with the biomarker but are not associated with the latent exposure. Based on our model introduced in the Subjects and Methods Section, under the standardized scale, $\sigma^2_{Y,j}$s are equal constants and thus can be cancelled out from the numerator and denominator of $\widehat{\theta}^{IVW}$. As GWAS sample sizes ($N_Y$ and $N_B$) go to infinity, we can derive that

$$\begin{aligned}\widehat{\theta}^{IVW} &= \frac{\sum_{j=1}^{M} \frac{\widehat{\beta}_{Y,j}\widehat{\beta}_{B,j}}{\sigma^2_{Y,j}}}{\sum_{j=1}^{M} \frac{\widehat{\beta}^2_{B,j}}{\sigma^2_{Y,j}}} \\ &= \frac{\sum_{j=1}^{p_v M} \frac{\widehat{\beta}_{Y,j}\widehat{\beta}_{B,j}}{\sigma^2_{Y,j}} + \sum_{j=p_v M+1}^{M} \frac{\widehat{\beta}_{Y,j}\widehat{\beta}_{B,j}}{\sigma^2_{Y,j}}}{\sum_{j=1}^{p_v M} \frac{\widehat{\beta}^2_{B,j}}{\sigma^2_{Y,j}} + \sum_{j=p_v M+1}^{M} \frac{\widehat{\beta}^2_{B,j}}{\sigma^2_{Y,j}}}\end{aligned}$$

For the first $p_v M$ valid IVs, we have

$$\begin{aligned}\mathrm{E}\left(\widehat{\beta}_{Y,j}\widehat{\beta}_{B,j}\right) &= \mathrm{E}\left[\mathrm{E}\left(\widehat{\beta}_{Y,j}\widehat{\beta}_{B,j}\right)|\beta_{x,j},\gamma_{B,j},\theta,\theta_B\right] \\ &= \mathrm{E}\left[cov\left(\widehat{\beta}_{Y,j},\widehat{\beta}_{B,j}\right) + \mathrm{E}\left(\widehat{\beta}_{Y,j}\right)\mathrm{E}\left(\widehat{\beta}_{B,j}\right)|\beta_{x,j},\gamma_{B,j},\theta,\theta_B\right] \\ &= \mathrm{E}\left[\theta\beta_{x,j}(\theta_{B,j}\beta_{x,j} + \gamma_{B,j})\right] \\ &= \theta\theta_B h_x^2, \text{ for } j = 1,\ldots,p_v M,\end{aligned}$$

where $\gamma_{B,j}$ and $\theta_B$ correspond to $\gamma_{k,j}$ and $\theta_k$, respectively, in our original model in the single-biomarker case. For the rest of the $(1-p_v)M$ invalid IVs, $\beta_{x,j} = 0$, and thus we have $\mathrm{E}\left(\widehat{\beta}_{Y,j}\widehat{\beta}_{B,j}\right) = 0$, for $j = p_vM+1,\ldots,M$.

Similarly, we can derive that $\mathrm{E}(\widehat{\beta}^2_{B,j}) = \theta_B^2 h_x^2 + h^2_{\gamma_B} + \sigma^2_{B,j}$ for $j = 1,\ldots,p_vM$, and for $j = p_vM + 1,\ldots,M$ we have $\mathrm{E}(\widehat{\beta}^2_{B,j}) = h^2_{\gamma_B} + \sigma^2_{B,j}$. Given that $\sigma^2_{B,j} \to 0$ as $N_B \to \infty$, we have

$$
\begin{aligned}
\widehat{\theta}^{IVW} &\xrightarrow[N_Y,N_B\to\infty]{p} \frac{p_vM\theta\theta_B h_x^2}{p_vM\left(\theta_B^2 h_x^2 + h^2_{\gamma_B}\right) + (1-p_v)Mh^2_{\gamma_B}} \\
&\xrightarrow[N_Y,N_B\to\infty]{p} \frac{p_v\theta\theta_B h_x^2}{p_v\theta_B^2 h_x^2 + h_\gamma^2},
\end{aligned}
$$

which leads to an asymptotic bias, $\theta\left[p_v\theta_B h_x^2/\left(p_v\theta_B^2 h_x^2 + h_\gamma^2\right) - 1\right]$.

### Appendix B: Derivation of the Estimating Equations in MRLE

Recall the assumed random effects model:

$$
\begin{aligned}
X &= \sum_{j=1}^{M}\beta_{x,j}G_j + \epsilon_x, \\
Y &= \theta X + \epsilon_y, \\
B_k &= \theta_k X + \sum_{j=1}^{M}\gamma_{k,j}G_j + \epsilon_{B_k},\ k = 1,2,\ldots,K,
\end{aligned}
$$

where $X, Y, B_k$s and $G_j$s denote the latent exposure, the outcome, the observable traits that are co-regulated by $X$ (we will use “biomarkers” as an example in the following text), and the SNPs that are selected as IVs for the MR analysis. Detailed model assumptions are described in Section 2 of the main manuscript.

The proposed MRLE method requires only the summary-level association statistics of $G_j$s on $B_k$s and $Y$, which we denote as $\{\widehat{\beta}_{B_k,j}, \sigma^2_{B_k,j}\}$, $k = 1,2,\ldots,K$, and $\{\widehat{\beta}_{Y,j}, \sigma^2_{Y,j}\}$, respectively, where $\widehat{\beta}$ and $\sigma^2$ refer to the estimated association coefficient and its standard error, respectively. From the model we can obtain the observable models:

$$
\begin{aligned}
B_k &= \sum_{j=1}^{M} (\theta_k \beta_{x,j} + \gamma_{k,j}) G_j + (\theta_k \epsilon_x + \epsilon_{B_k}), \; k = 1, 2, \ldots, K, \\
Y &= \sum_{j=1}^{M} \theta \beta_{x,j} G_j + (\theta \epsilon_x + \epsilon_y),
\end{aligned}
$$

from which we see that $\widehat{\beta}_{B_k,j}$ has a mean of $\theta_k \beta_{x,j} + \gamma_{k,j}$, which is the estimated total effect of $G_j$ on $B_k$, and a variance of $\sigma^2_{B_k,j}$, and similarly, $\widehat{\beta}_{Y,j}$ has a mean of $\theta\beta_{x,j}$, which is the total effect of $G_j$ on $B_k$, and a variance of $\sigma^2_{Y,j}$.

Given the prior assumption that $\beta_{x,j}$s are independent and identically (i.i.d.) distributed with mean 0 and variance $h_x^2$, $\gamma_{k,j}$s are i.i.d. distributed with mean 0 and variance $h^2_{\gamma_k}$, and the independence assumption between $\beta_{x,j}$s and $\gamma_{k,j}$s, we can derive the second-order moments of $\{\widehat{\beta}_{Y,j}, \widehat{\beta}_{B_1,j}, \ldots, \widehat{\beta}_{B_K,j}, j = 1, 2, \ldots, M\}$,

$$
\begin{aligned}
\mathrm{E}(\widehat{\beta}_{Y,j}\widehat{\beta}_{B_k,j}) &= \mathrm{E}[\mathrm{E}(\widehat{\beta}_{Y,j}\widehat{\beta}_{B_k,j})|\beta_{x,j}, \gamma_{k,j}] \\
&= \mathrm{E}[cov(\widehat{\beta}_{Y,j}, \widehat{\beta}_{B_k,j})|\beta_{x,j}, \gamma_{k,j}] + \mathrm{E}[\mathrm{E}(\widehat{\beta}_{Y,j})\mathrm{E}(\widehat{\beta}_{B_k,j})|\beta_{x,j}, \gamma_{k,j}] \\
&= \mathrm{E}[\theta\beta_{x,j}(\theta_k\beta_{x,j} + \gamma_{k,j})|\beta_{x,j}, \gamma_{k,j}] \\
&= \theta\theta_k \mathrm{E}(\beta^2_{x,j}) \\
&= \theta\theta_k h_x^2, \;\; 1 \leqslant k \leqslant K, \\
\mathrm{E}(\widehat{\beta}^2_{B_k,j}) &= \mathrm{E}[\mathrm{E}(\widehat{\beta}^2_{B_k,j})|\beta_{x,j}, \gamma_{k,j}] \\
&= \mathrm{E}[var(\widehat{\beta}_{B_k,j})|\beta_{x,j}, \gamma_{k,j}] + \mathrm{E}[(\mathrm{E}(\widehat{\beta}_{B_k,j}))^2|\beta_{x,j}, \gamma_{k,j}] \\
&= \sigma^2_{B_k,j} + \mathrm{E}[(\theta_k\beta_{x,j} + \gamma_{k,j})^2|\beta_{x,j}, \gamma_{k,j}] \\
&= \theta_k^2 E(\beta^2_{x,j}) + \mathrm{E}(\gamma^2_{k,j}) + \sigma^2_{B_k,j} \\
&= \theta_k^2 h_x^2 + h^2_{\gamma_k} + \sigma^2_{B_k,j}, \;\; 1 \leqslant k \leqslant K,
\end{aligned}
$$

$$
\begin{aligned}
\mathrm{E}(\widehat{\beta}_{B_k,j}\widehat{\beta}_{B_{k'},j}) &= \mathrm{E}[\mathrm{E}(\widehat{\beta}_{B_k,j}\widehat{\beta}_{B'_k,j})|\beta_{x,j},\gamma_{k,j},\gamma_{k',j}] \\
&= \mathrm{E}[cov(\widehat{\beta}_{B_k,j},\widehat{\beta}_{B'_k,j})|\beta_{x,j},\gamma_{k,j},\gamma_{k',j}] + \mathrm{E}[\mathrm{E}(\widehat{\beta}_{B_k,j})\mathrm{E}(\widehat{\beta}_{B'_k,j})|\beta_{x,j},\gamma_{k,j},\gamma_{k',j}] \\
&= c_{k,k'} + \mathrm{E}[(\theta_k\beta_{x,j}+\gamma_{k,j})(\theta_{k'}\beta_{x,j}+\gamma_{k',j})|\beta_{x,j},\gamma_{k,j},\gamma_{k',j}] \\
&= \theta_k\theta_{k'}\mathrm{E}(\beta_{x,j}^2) + c_{k,k'} \\
&= \theta_k\theta_{k'}h_x^2 + c_{k,k'},\quad 1 \leqslant k < k' \leqslant K,
\end{aligned}
$$

where $c_{k,k'}$ denotes the covariance of $\widehat{\beta}_{B_k,j}$ and $\widehat{\beta}_{B_{k'},j}$ that is commonly observed due to sample overlap between GWASs of correlated biomarkers.

If the summary-level data for biomarkers $B_k$ and $B_{k'}$ are from two overlapping GWASs (i.e. there is overlapping individuals in the GWASs), then $c_{k,k'}$s can be estimated by fitting a bivariate LD score regression.[2] Suppose that we have the summary-level estimate for the effect of $G_j$s on $B_k$ and $B_{k'}$, which we denote as $\{\widehat{\beta}_{k,j}, j = 1,2,\ldots,M\}$ and $\{\widehat{\beta}_{k',j}, j = 1,2,\ldots,M\}$, respectively. We assume that $\widehat{\beta}_{k,j}$ is observed on the standardized scale, i.e. $var(B_k) = 1$, so that $var(\widehat{\beta}_{k,j}) = 1/n_{k,j}$, where $n_{k,j}$ denotes the sample size for $G_j$ in the GWAS for $B_k$. We further denote the true joint effect size and marginal effect size as $\beta_{k,j}$s and $\tilde{\beta}_{k,j}$s, respectively, the genetic covariance between $B_k$ and $B_{k'}$ as $h_{k,k'} = cov(\beta_{k,j},\beta_{k',j})$, and the LD score of $G_j$ as $l_j = \sum_{j^*=1}^{M} r_{j,j^*}^2$, $j = 1,2,\ldots,M$, where $r_{j,j^*}$ denotes the correlation between $G_j$ and $G_{j^*}$. Given $\mathrm{E}(\widehat{\beta}_{k,j}) = \tilde{\beta}_{k,j}$, $\tilde{\beta}_{k,j} = \sum_{j^*=1}^{M} r_{j,j^*}\beta_{k,j^*}$, $j = 1,2,\ldots,M$, we then have

$$
\begin{aligned}
\mathrm{E}(\widehat{\beta}_{k,j},\widehat{\beta}_{k,j}|l_j) &= \mathrm{E}[(\widehat{\beta}_{k,j}-\tilde{\beta}_{k,j}+\tilde{\beta}_{k,j})(\widehat{\beta}_{k',j}-\tilde{\beta}_{k',j}+\tilde{\beta}_{k',j})|l_j] \\
&= \mathrm{E}[(\widehat{\beta}_{k,j}-\tilde{\beta}_{k,j})(\widehat{\beta}_{k',j}-\tilde{\beta}_{k',j})|l_j] + \mathrm{E}[(\widehat{\beta}_{k,j}-\tilde{\beta}_{k,j})\tilde{\beta}_{k',j}|l_j] \\
&+ \mathrm{E}[\tilde{\beta}_{k,j}(\widehat{\beta}_{k',j}-\tilde{\beta}_{k',j})|l_j] + \mathrm{E}[\tilde{\beta}_{k,j}\tilde{\beta}_{k',j}|l_j] \\
&= c_{k,k'} + \mathrm{E}[(\sum_{j^*=1}^{M} r_{j,j^*}\beta_{k,j^*})(\sum_{j^*=1}^{M} r_{j,j^*}\beta_{k',j^*})|l_j] \\
&= c_{k,k'} + \sum_{j_1=1}^{M}\sum_{j_2=1}^{M} r_{j,j_1}r_{j,j_2}\mathrm{E}(\beta_{k,j_1}\beta_{k',j_2}) \\
&= c_{k,k'} + \sum_{j^*=1}^{M} r_{j,j^*}^2 E(\beta_{k,j^*}\beta_{k',j^*}) \\
&= c_{k,k'} + h_{k,k'}l_j,
\end{aligned}
$$

which gives

$$\mathrm{E}(z_{k,j}z_{k',j}) = \sqrt{n_{k,j}n_{k',j}}c_{k,k'} + \sqrt{n_{k,j}n_{k',j}}h_{k,k'}l_j,$$

where $z_{k,j} = \sqrt{n_{k,j}}\widehat{\beta}_{k,j}$ denotes the summary-level z-score. We can see that $c_{k,k'}$ can be estimated by obtaining the intercept of an LD score regression of $z_{k,j}z_{k',j}$s on the LD score $l_j$s. We notice that in the second-order moments, the per-SNP genetic heritability of the latent exposure, $h_x^2$, always shows up alongside of $\theta_a\theta_b$ where $\theta_a, \theta_b \in \{\theta, \theta_1, \ldots, \theta_K\}$. To avoid identifiability issue, we reparameterize the model with $\mu = \theta h_x$, $\mu_k = \theta_k h_x$, $k = 1, 2, \ldots, K$, and denote the vector of the unknown parameters as $\eta = (\mu, \mu_1, \ldots, \mu_K, h_{\gamma_1}^2, \ldots, h_{\gamma_K}^2)^T$.

Finally, we construct the estimating equations by equating the second-order moments to the corresponding sample moments

$$\Psi(X^s, \eta) = (1/M)\sum_{j=1}^{M}\psi(X_j^s, \eta) = 0,$$

where $\psi(X_j^s, \eta) = \Big(\psi_1(X_j^s, \eta), \psi_2(X_j^s, \eta), \ldots, \psi_d(X_j^s, \eta)\Big)^T$, with

$$\psi_k(X_j^s, \eta) = \widehat{\beta}_{Y,j}\widehat{\beta}_{B_k,j} - \mu_k\mu, \;\; 1 \leqslant k \leqslant K,$$

$$\psi_{K+k}(X_j^s, \eta) = \widehat{\beta}_{B_k,j}^2 - \sigma_{B_k,j}^2 - h_{\gamma_k}^2 - \mu_k^2, \;\; 1 \leqslant k \leqslant K,$$

$$\psi_{\nu(k,k')}(X_j^s, \eta) = \widehat{\beta}_{B_k,j}\widehat{\beta}_{B_{k'},j} - c_{k,k'} - \mu_k\mu_{k'}, \;\; 1 \leqslant k < k' \leqslant K.$$

More specifically, we have a set of estimating equations,

$$\begin{aligned}
\frac{1}{M}\sum_{j=1}^{M}\widehat{\beta}_{Y,j}\widehat{\beta}_{B_k,j} &= \mu_k\mu, \;\; 1 \leqslant k \leqslant K,\\
\frac{1}{M}\sum_{j=1}^{M}\widehat{\beta}_{B_k,j}^2 &= \sigma_{B_k,j}^2 + h_{\gamma_k}^2 + \mu_k^2, \;\; 1 \leqslant k \leqslant K,\\
\frac{1}{M}\sum_{j=1}^{M}\widehat{\beta}_{B_k,j}\widehat{\beta}_{B_{k'},j} &= c_{k,k'} + \mu_k\mu_{k'}, \;\; 1 \leqslant k < k' \leqslant K,
\end{aligned}$$

Recall that the generalized method of moments (GMM) estimator of the vector of model parameters, $\eta = (\mu, \mu_1, \ldots, \mu_K, h_{\gamma_1}^2, \ldots, h_{\gamma_K}^2)^T$, is defined as

$$\widehat{\eta}^{\mathrm{GMM}} = \underset{\eta}{\operatorname{argmin}}\, Q_{\widehat{W}}(\eta),$$

which is the global minimizer of the objective function $Q_{\widehat{W}}(\eta) = \Psi(X^s, \eta)^T \widehat{W} \Psi(X^s, \eta)$, where $\widehat{W}$ is a user-defined positive semi-definite weighting matrix. Here we consider the optimal choice of $\widehat{W}$ with minimum asymptotic covariance, $\widehat{W}^{\text{opt}}$, which is the inverse of covariance matrix of the estimating equations $\Omega(\eta) = \mathrm{E}\left(\psi(X^s, \eta_0)\psi(X^s, \eta_0)^T\right)$, based on the true parameter value $\eta_0$.

We now derive the closed form of $\Omega(\eta)$. To simplify notations, we denote $\psi_i(X^s, \eta)$ by $\psi_i$, $i = 1, 2, \ldots, d$. The $(i, j)$-th and the $(j, i)$-th entry of $\Omega(\eta)$ are both equal to

$$
\begin{aligned}
E(\psi_i \psi_j) &= E\left[E\left(\psi_i \psi_j\right) | \beta_x, \gamma_1, \ldots, \gamma_K\right] \\
&= E\left[E\left(\psi_i\right) E\left(\psi_j\right) + cov\left(\psi_i, \psi_j\right) | \beta_x, \gamma_1, \ldots, \gamma_K.\right]
\end{aligned}
$$

We can then derive the following results:

$$
\begin{aligned}
\mathrm{E}(\psi_k^2) &= \theta^2 h_x^2(2\theta_k^2 h_x^2 + h_k^2) + \theta^2 h_x^2\sigma_k^2 + (\theta_k^2 h_x^2 + h_k^2)\sigma_Y^2 + 2\theta\theta_k h_x^2 c_{y,k} + \sigma_k^2\sigma_Y^2 + c_{y,k}^2,\\
\mathrm{E}(\psi_k\psi_l) &= 2\theta_k\theta_l\theta^2 h_x^4 + \theta_k\theta_l h_x^2\sigma_Y^2 + \theta\theta_k h_x^2 c_{y,l} + \theta^2 h_x^2 c_{k,l} + \theta\theta_l h_x^2 c_{y,k}\\
&+ \sigma_y^2 c_{k,l} + c_{y,k}c_{y,l},\\
\mathrm{E}(\psi_{K+k}\psi_k) &= 2\theta\theta_k h_x^2(\theta_k^2 h_x^2 + h_k^2 + \sigma_k^2) + 2c_{y,k}(\theta_k^2 h_x^2 + h_k^2 + \sigma_k^2),\\
\mathrm{E}(\psi_{K+k}\psi_l) &= 2\theta\theta_l\theta_k^2 h_x^4 + 2\theta_k\theta_l h_x^2 c_{y,k} + 2\theta\theta_k h_x^2 c_{k,l} + 2c_{k,l}c_{y,k},\\
\mathrm{E}(\psi_{\nu(k,k')}\psi_k) &= \theta\theta_{k'} h_x^2(2\theta_k^2 h_x^2 + h_k^2) + \theta_{k'}\theta h_x^2\sigma_k^2 + \theta_k\theta_{k'} h_x^2 c_{y,k}\\
&+ (\theta_k^2 h_x^2 + h_k^2)c_{y,k'} + \theta\theta_k h_x^2 c_{k,k'} + c_{y,k'}\sigma_k^2 + c_{k,k'}c_{y,k},\\
\mathrm{E}(\psi_{\nu(k,k')}\psi_l) &= 2\theta\theta_k\theta_{k'}\theta_l h_x^4 + \theta_{k'}\theta_l h_x^2 c_{y,k} + \theta_{k'}\theta h_x^2 c_{k,l} + \theta_k\theta_l h_x^2 c_{y,k'}\\
&+ \theta\theta_k h_x^2 c_{k',l} + c_{k',l}c_{y,k} + c_{y,k'}c_{k,l},\\
\mathrm{E}(\psi_{K+k}^2) &= 2[(\theta_k^2 h_x^2 + h_k^2)^2 + 2\sigma_k^2(\theta_k^2 h_x^2 + h_k^2)] + 2\sigma_k^4,\\
\mathrm{E}(\psi_{K+k}\psi_{K+l}) &= 2\theta_k^2\theta_l^2 h_x^4 + 4\theta_k\theta_l h_x^2 c_{k,l},\\
\mathrm{E}(\psi_{\nu(k,k')}\psi_{K+k}) &= 2\theta_k\theta_{k'} h_x^2(h_k^2 + \theta_k^2 h_x^2) + 2\theta_k\theta_{k'} h_x^2\sigma_k^2 + 2(\theta_k^2 h_x^2 + h_k^2)c_{k,k'} + 2\sigma_k^2 c_{k,k'},\\
\mathrm{E}(\psi_{\nu(k,k')}\psi_{K+l}) &= 2\theta_k\theta_{k'}\theta_l^2 h_x^4 + 2\theta_{k'}\theta_l h_x^2 c_{k,l} + 2\theta_k\theta_l h_x^2 c_{k',l} + 2c_{k,l}c_{k',l},\\
\mathrm{E}(\psi_{\nu(k,k')}^2) &= 2\theta_k^2\theta_{k'}^2 h_x^4 + (\theta_{k'}^2 h_x^2 + h_{k'}^2)\sigma_k^2 + (\theta_k^2 h_x^2 + h_k^2)\sigma_{k'}^2 + 2\theta_k\theta_{k'} h_x^2 c_{k,k'}\\
&+ (\theta_k^2 h_{k'}^2 + \theta_{k'}^2 h_k^2)h_x^2 + h_k^2 h_{k'}^2 + \sigma_k^2\sigma_{k'}^2 + c_{k,k'}^2,\\
\mathrm{E}(\psi_{\nu(k,l_1)}\psi_{\nu(k,l_2)}) &= \theta_{l_1}\theta_{l_2} h_x^2(2h_x^2\theta_k^2 + h_k^2) + \theta_{l_1}\theta_{l_2} h_x^2\sigma_k^2 + \theta_{l_1}\theta_k h_x^2 c_{k,l_2} + \theta_k\theta_{l_2} h_x^2 c_{k,l_1}\\
&+ (\theta_k^2 h_x^2 + h_k^2)c_{l_1,l_2} + \sigma_k^2 c_{l_1,l_2} + c_{k,l_1}c_{k,l_2}
\end{aligned}
$$

$$
\begin{aligned}
\mathrm{E}(\psi_{\nu(k,l_1)}\psi_{\nu(l_1,l_2)}) &= \theta_k\theta_{l_2}h_x^2(2h_x^2\theta_{l_1}^2 + h_{l_1}^2) + \theta_k\theta_{l_2}h_x^2\sigma_{l_1}^2 + \theta_k\theta_{l_1}h_x^2 c_{l_1,l_2} + \theta_{l_1}\theta_{l_2}h_x^2 c_{k,l_1} \\
&\quad + (\theta_{l_1}^2 h_x^2 + h_{l_1}^2)c_{k,l_2} + \sigma_{l_1}^2 c_{k,l_2} + c_{k,l_1}c_{l_1,l_2}, \\
\mathrm{E}(\psi_{\nu(k,l)}\psi_{\nu(k',l)}) &= \theta_k\theta_{k'}h_x^2(2h_x^2\theta_l^2 + h_l^2) + \theta_k\theta_{k'}h_x^2\sigma_l^2 + \theta_k\theta_l h_x^2 c_{l,k'} + \theta_l\theta_{k'}h_x^2 c_{k,l} \\
&\quad + (\theta_l^2 h_x^2 + h_l^2)c_{k,k'} + \sigma_l^2 c_{k,k'} + c_{l,k}c_{l,k'}, \\
\mathrm{E}(\psi_{\nu(k,l_1)}\psi_{\nu(k',l_2)}) &= 2\theta_k\theta_{k'}\theta_{l_1}\theta_{l_2}h_x^4 + \theta_{l_1}\theta_{l_2}h_x^2 c_{k,k'} + \theta_{l_1}\theta_{k'}h_x^2 c_{k,l_2} + \theta_k\theta_{l_2}h_x^2 c_{l_1,k'} \\
&\quad + \theta_k\theta_{k'}h_x^2 c_{l_1,l_2} + c_{k,k'}c_{l_1,l_2} + c_{k,l_2}c_{k',l_1},
\end{aligned}
$$

where $\nu(k,k') = 2K + \sum_{l=1}^{k-1}(K-l) + (k'-k)$, $\sigma_Y^2 = \theta^2\sigma_x^2/n_y$, and $\sigma_k^2 = (\theta_k^2\sigma_x^2 + \sigma_{\epsilon_k}^2)/n_y$, $k = 1, 2\ldots, K$. Since $\widehat{W}^{\text{opt}}$ is a function of the unknown true parameter value $\eta_0$, we conduct inference using an iterative two-step GMM algorithm, which will be described in Appendix C.

Assume that $\widehat{W}$ converges in probability ($\xrightarrow{p}$) to a positive semi-definite matrix $W$, and denote the true parameter value as $\eta_0$. The consistency of the GMM estimator $\widehat{\eta}^{\text{GMM}}$ is guaranteed given the following assumptions: (A.1) $W\,\mathrm{E}\left[\psi(X^s,\eta)\right] = 0$ if and only if $\eta = \eta_0$; (A.2) the parameter space $\Theta \subset \mathbb{R}^p$ is compact; (A.3) $\psi(X^s,\eta)$ is continuous at each $\eta \in \Theta$ with probability one; and (A.4) $\mathrm{E}\left[\sup_{\eta\in\Theta}\|\psi(X^s,\eta)\|\right] < \infty$, where $\|\cdot\|$ denotes the $L^2$ norm.[3,4,5] To construct a variance estimator for $\widehat{\eta}^{\text{GMM}}$, we first introduce several additional terms including the $d \times d$ covariance matrix of the estimating equations, $\Omega(\eta) = \mathrm{E}_{X^s}\left[\Psi(X^s,\eta)\Psi(X^s,\eta)^T\right]$, and the expectation of the $d \times p$ derivative matrix of the estimating equations, $D(\eta) = \mathrm{E}_{X^s}\left[\nabla_\eta\Psi(X^s,\eta)\right]$. We will denote $\Omega(\eta_0)$ as $\Omega_0$ and $D(\eta_0)$ as $D_0$ for simplicity. Given that $\widehat{\eta}^{\text{GMM}}$ is consistent, $\sqrt{M}\left(\widehat{\eta}^{\text{GMM}} - \eta_0\right) \xrightarrow{d} \mathcal{N}(0, (D_0^T D_0)^{-1}D_0^T W\Omega_0 W^T D_0(D_0^T D_0)^{-1})$, where $\xrightarrow{d}$ denotes convergence in distribution (weak convergence), holds under the following conditions: (B.1) $\psi(X^s,\eta)$ is continuously differentiable in a neighborhood of $\eta_0$, $\mathcal{U}(\eta_0)$, with probability one; (B.2) $\mathrm{E}_{X^s}[\psi(X^s,\eta)^2] < \infty, \forall\eta \in \Theta$; (B.3) $\mathrm{E}[\sup_{\eta\in\mathcal{U}(\eta_0)}\|\nabla_\eta g(Y_t,\eta)\|] < \infty$; and (B.4) $D_0^T W D_0$ is invertible.[3,4,5] The asymptotic theory of GMM further suggests that an optimal choice for $W$ is a matrix proportional to $\Omega_0^{-1}$, which gives an optimal asymptotic covariance matrix $(D_0^T\Omega_0^{-1}D_0)^{-1}$.

### APPENDIX C: IMPLEMENTATION OF MRLE USING REGULARIZED NEWTON-RAPHSON'S METHOD

The GMM estimator of the vector of unknown parameters $\eta$ is defined as $\widehat{\eta}^{\text{GMM}} = \text{argmin}_\eta\, Q_{\widehat{W}}(\eta)$, where $Q_{\widehat{W}}(\eta) = \Psi(X^s, \eta)^T \widehat{W} \Psi(X^s, \eta)$ is the objective function, and $\widehat{W}$ is a positive semidefinite weighting matrix that is typically specified based on the summary data $X^s$. $\widehat{\eta}^{\text{GMM}}$ can be obtained by solving the equation, $D(\eta)\widehat{W}\Psi(\eta) = 0$, where $D(\eta) = \text{E}\left(\nabla_\eta \Psi(X^s, \eta)\right)$ is the expectation of the $d \times p$ derivative matrix of the estimating equations $\Psi(X^s, \eta)$, $T(\eta) = D(\eta)\widehat{W}\Psi(\eta)$ is the gradient of the objective function $Q_{\widehat{W}}(\eta)$.

We implement the two-step GMM algorithm.[3] In step one, we set the weighting matrix to $\widehat{W} = I$ and compute a preliminary GMM estimate, $\widehat{\eta}_{(0)}$. In step two, we set $\widehat{W} = \left(\frac{1}{M}\sum_{j=1}^{M} \psi(x_j^s, \hat{\eta}_{(0)})\psi(x_j^s, \hat{\eta}_{(0)})^T\right)^{-1}$, which is the sample estimate of the optimal weighting matrix $\widehat{W}^{\text{opt}}$, and compute the final GMM estimator, $\widehat{\eta}^{\text{GMM}}$. It is easy to see that $\widehat{W} \xrightarrow{p} \Omega^{-1}$, and thus the resulting estimator is asymptotically the most efficient among the asymptotically normal estimators (Hanson, 1982). To compute the GMM estimator in each of the two steps, we adopt a regularized Newton Raphson (NR) algorithm[6,7] which proceeds as follows.

1. We first generate $\theta^{(0)}, \theta^{(0,*)} \sim \text{U}(-\sqrt{0.1}, \sqrt{0.1})$, $\theta_k^{(0)}\theta_k^{(0,*)} \sim \text{U}(0, \sqrt{0.7})$, $h_x^{2\,(0)} = \left((1/M)\sum_{j=1}^{M} \widehat{\beta}_{Y,j}^2 - \widehat{\sigma^2}_{Y,j}\right)/\theta^{(0,*)2}$, $h_{\gamma_k}^{2\,(0)} = (1/M)\sum_{j=1}^{M} \widehat{\beta}_{B_k,j}^2 - \widehat{\sigma^2}_{B_k,j} - h_x^{2\,(0)}\theta_k^{(0,*)2}$, then set $\mu^{(0)} = \theta^{(0)} h_x^{(0)}$, and $\mu_k^{(0)} = \theta_k^{(0)} h_x^{(0)}$, which gives a set of initial values, $\widehat{\eta}^{(0)} = \left(\mu^{(0)}, \{\mu_k^{(0)}\}_{k=1}^K, \{h_{\gamma_k}^{2\,(0)}\}_{k=1}^K\right)^T$.

2. At the $(t+1)$-th iteration, update $\widehat{\eta}^{(t+1)} = \widehat{\eta}^{(t)} - \left(J(\widehat{\eta}^{(t)}) + \mu_t I\right)^{-1} T(\widehat{\eta}^{(t)})$, where $J(\widehat{\eta})$ denotes the Hessian matrix of $Q_{\widehat{W}}(\eta)$,
$$J(\widehat{\eta}) = D^T(\eta)\widehat{W}D(\eta) + \begin{bmatrix} \Psi^T(\eta)\widehat{W}\frac{\partial}{\partial\eta}D_1(\eta) \\ \cdots \\ \Psi^T(\eta)\widehat{W}\frac{\partial}{\partial\eta}D_p(\eta) \end{bmatrix},$$
$D_i(\eta)$ denotes the $i$-th column of $D(\eta)$, $i = 1, 2, \ldots, p$, and $\mu_t = \|T(\widehat{\eta}^{(t)})\|_2$ is a small value that is updated iteratively and added to the diagonal entries of $J(\eta)$ to improve convergence near the potential singular solutions.

3. Repeat step 2 until a pre-specified convergence criterion (e.g. $\|\widehat{\eta}^{(t)} - \widehat{\eta}^{(t-1)}\| \leqslant \varepsilon$ where $\varepsilon$ is a small-valued term) is met.

4. Implement steps 1 to 3 with multiple initial values to obtain multiple solutions, remove the ones that have at least two of the following entries that fall out of their ranges, $\theta_k \in (0, \infty)$, $h^2_{\gamma_k} \in (0, 1/M)$, and obtain $N$ candidate solutions $\{\widehat{\eta}^{[1]}, \ldots, \widehat{\eta}^{[N]}\}$. We then obtain the GMM estimator, $\widehat{\eta}^{\text{GMM}} = \text{argmin}_i\, Q_{\widehat{W}}(\widehat{\eta}^{[i]})$.

The expectation of the $d \times p$ derivative matrix of the estimating equations $\Psi(X^s, \eta)$ is

$$
\begin{aligned}
D(\eta) = &\ \mathrm{E}\left[\nabla_\eta \Psi(X^s, \eta)\right] \\
= &\begin{bmatrix}
\mu_1 & \mu\ \ldots\ 0 & 0\ \ldots\ 0 \\
\ldots & \ldots & \ldots \\
\mu_K & 0\ \ldots\ \mu & 0\ \ldots\ 0 \\
0 & 2\mu_1\ \ldots\ 0 & 1\ \ldots\ 0 \\
\ldots & \ldots & \ldots \\
0 & 0\ \ldots\ 2\mu_K & 0\ \ldots\ 1 \\
0 & \mu_2\ \mu_1\ \ldots\ 0 & 0\ \ldots\ 0 \\
\ldots & \ldots & \ldots \\
0 & 0 \ldots \mu_{k'} \ldots \mu_k \ldots 0 & 0\ \ldots\ 0 \\
\ldots & \ldots & \ldots \\
0 & 0\ \ldots\ \mu_K\ \ \mu_{K-1} & 0\ \ldots\ 0
\end{bmatrix}_{d \times p}.
\end{aligned}
$$

Let $D = [D_1, \ldots, D_{2(K+1)}]$. The $(i, j)$-th entry of $\nabla_\eta D_1$ is

$$
\left(\nabla_\eta D_1\right)_{i,j} = I(j = i + 1, i \in \{1, \ldots, K\}).\ i = 1, \ldots, d, j = 1, \ldots, K,
$$

For $k = 1, \ldots, K$, $\left(\nabla_\eta D_{k+1}\right)_{i,j} = 0$ holds for all $i = 1, \ldots, d,\ j = 1, \ldots, p$, except that $\left(\nabla_\eta D_{k+1}\right)_{K+k,k+1} = 2$, and when $k > 1$, for $j < k$, the $(2K + \sum_{l=1}^{j-1}(K-l) + (k-j), 1+j)$-th entry of $\nabla_\eta D_{k+1}$ is 1, and for $j > k$, the $(2K + \sum_{l=1}^{k-1}(K-l) + (j-k), 1+j)$-th entry of $\nabla_\eta D_{k+1}$ is 1. Additionally, we have $\nabla_\eta D_{K+k+1} = \mathbf{0}_{d \times p}$.

### Appendix D: Detailed Simulation Settings

#### *D.1. A pilot study to compare individual-level and summary-level simulations*

We first conducted individual-level data-based simulations. Specifically, we set the GWAS sample size to $N = 6 \times 10^4$, and assumed there were a total of $K = 6$ biomarkers that were co-regulated by the latent exposure. We started with collecting genotype data for $N$ individuals that were randomly selected from the approximately 330K unrelated individuals of European origin from UK Biobank.[8] We first conducted LD pruning to select $M = 126,627$ common SNPs, i.e., SNPs with MAF>0.01, that had an absolute pairwise correlation lower than 0.1 within a 500kb genetic distance. These SNPs were then included in our analysis. Instead of assuming all SNPs to be causal as in our "exposure" and "biomarker" model, we considered a more realistic setting where only a small proportion of the SNPs, which was set to 1% here, are causal, i.e., having non-zero effects on the trait (biomarker or exposure). We set $\theta_k$ to $\sqrt{0.3}$ for $k = 1, 2, \ldots, K$, so that $X$ explained 30% of the variability of each biomarker. We further set the total heritability of each biomarker ($H^2_{\text{B.total}}$) to 0.2, and the proportion explained by the association with $X$ to 0.2, resulting in a total heritability of each biomarker explained by $X$ ($H^2_{B.X}$) between 0.04 and 0.09. The effect of the exposure $X$ on the outcome $Y$ ($\theta$) was set to either 0 (to evaluate type I error control) or 0.1 (to evaluate power).

We first selected a random 1% of the $M$ SNPs to be causal for the exposure and each biomarker. Specifically, we randomly selected $M_x = 0.01M$ SNPs to be truly associated with the exposure $X$, with effect sizes $\beta_{x,j}$s generated from $N(0, h_x^2)$, where $h_x^2$ is the per-SNP heritability of $X$ that can be calculated based on $H^2_{\text{B.total}}$, $H^2_{\text{B.X}}$ and $\theta_k$. Similarly, we randomly selected $M_{B_k} = 0.01M$ SNPs to be associated with $B_k$, with effect sizes $\gamma_{k,j}$s generated from $N(0, h^2_{B_k})$, $k = 1, 2, \ldots, K$, where $h^2_{B_k}$ can be calculated based on $H^2_{\text{B.total}}$ and $H^2_{\text{B.X}}$. We assumed there were unmeasured confounders between the exposure and the biomarkers, between different biomarkers, between the biomarkers and the outcome, and between the exposure and the outcome, that are not associated with the selected IVs. Specifically, we assume a correlation of 0.3 between each pair of residual terms among $\epsilon_y$ ("outcome model"), $\epsilon_x$ ("exposure model"), and $\epsilon_{B_k}$s ("biomarker models"). To simplify the calculation, we assumed the genotype, the exposure, the biomarkers, and the outcome were all standardized to have a zero mean and a unit variance. Based on our model, we could then calculate the variance of the residual terms $\epsilon_y$, $\epsilon_x$, and $\epsilon_{B_k}$s, based on which we simulated data for the exposure, the biomarkers, and the outcome for the $N$ individuals. Finally, we conducted GWAS analysis on each biomarker and the outcome using one-SNP-at-a-time regressions. The various tests were then applied to the GWAS summary data.

For summary-level simulations, we applied the various tests on data simulated based on the exact same simulation settings. One thing to note is that we first conducted individual-level simulations, where we used the bivariate LD score regression to calculate sample covariance matrices for the biomarkers ($c_{k,k'}$s; we then conducted summary-level simulations, where we directly simulated GWAS summary data based on the covariance matrices ($c_{k,k'}$s estimated from the corresponding individual-level data-based simulation. We used this strategy to ensure a fair comparison between the two types of simulations.

#### *D.2. Simulating on the summary level*

We can directly simulate GWAS summary-level association statistics instead of individual-level data to reduce computation time.[2] Specifically, for SNP $j$, we first generated the true effect sizes, $\beta_{x,j}$ and $\gamma_{k,j}$, $k = 1, \ldots, K$, then simulated the corresponding summary-level estimates based on the following model,

$$\begin{pmatrix} \widehat{\beta}_{B_1,j} \\ \cdots \\ \widehat{\beta}_{B_K,j} \\ \widehat{\beta}_{Y,j} \end{pmatrix} = \begin{pmatrix} \theta_1 \beta_{x,j} + \gamma_{1,j} \\ \cdots \\ \theta_K \beta_x + \gamma_{K,j} \\ \theta \beta_x \end{pmatrix} + \boldsymbol{\epsilon}_j.$$

The error term $\boldsymbol{\epsilon}_j$ accounts for the variability in the estimates that resulted from within-study variance and potential between-study correlation due to sample overlap. Given independence assumption of the SNPs, we generated $\boldsymbol{\epsilon}_j$s independently from a multivariate normal distribution, $\mathcal{MVN}(\mathbf{0}, \boldsymbol{\Gamma})$, with covariance matrix

$$\boldsymbol{\Gamma} = \begin{pmatrix} \boldsymbol{\Gamma}_{\boldsymbol{B}} & \mathbf{0} \\ \mathbf{0} & N_Y^{-1} \end{pmatrix},$$

where $\boldsymbol{\Gamma}_{\boldsymbol{B}}$ is a $K \times K$ matrix with the $(k, l)$-th entry equal to $(\boldsymbol{\Gamma}_{\boldsymbol{B}})_{k,l} = \left(N_{B_k,B_l} / (N_{B_k} N_{B_l})\right) cov_{B_k,B_l}$, $N_a$ denotes the GWAS sample size for $a \in \{Y, B_k, k = 1, \ldots, K\}$, $N_{B_k,B_l}$ and $cov(B_k, B_l)$ denote the overlapping GWAS sample size and covariance, respectively, between $B_k$ and $B_l$, $k, l = 1, \ldots, K$.

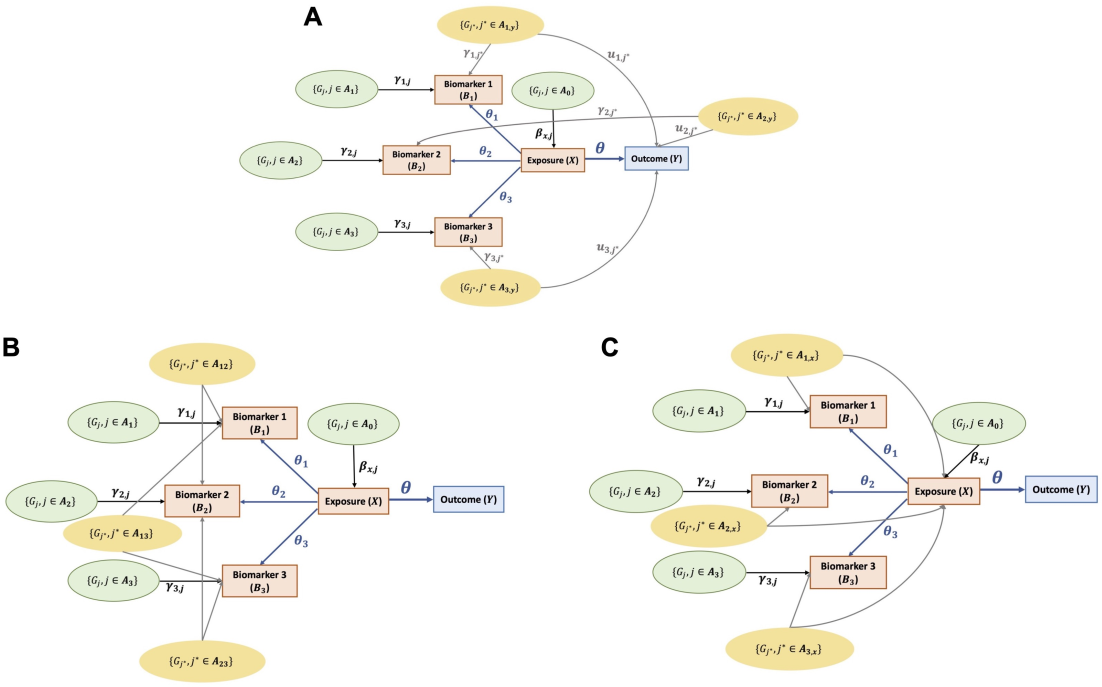


Figure S1. Causal paths between the SNPs ($G_j$s), the latent exposure of interest ($X$), the biomarkers co-regulated by the latent exposure ($B_k$s), and the outcome ($Y$) under three types of pleiotropy. The notations follow those in Figure 1. SNPs having various types of pleiotropic effects are highlighted in yellow ovals. **(A)** Pleiotropic effects between $B_k$s and $Y$: some SNPs $G_{j^*}$, $j^* \in \boldsymbol{A}_{k,y}$ have correlated effects on $Y$ ($u_{k,j^*}$s) and at least one of the $B_k$s ($\gamma_{k,j^*}$s). **(B)** Pleiotropic effects across $B_k$s: some SNPs $G_{j^*}$, $j^* \in \boldsymbol{A}_{k,k'}$ have correlated direct effects ($\gamma_{k,j^*}$ and $\gamma_{k',j^*}$) between at least two biomarkers $B_k$ and $B_{k'}$. **(C)** Pleiotropic effects between $B_k$s and $X$: some SNPs $G_{j^*}$, $j^* \in \boldsymbol{A}_{k,x}$ have correlated direct effects on $X$ ($\beta_{x,j^*}$) and at least one of the $B_k$s ($\gamma_{k,j^*}$s).

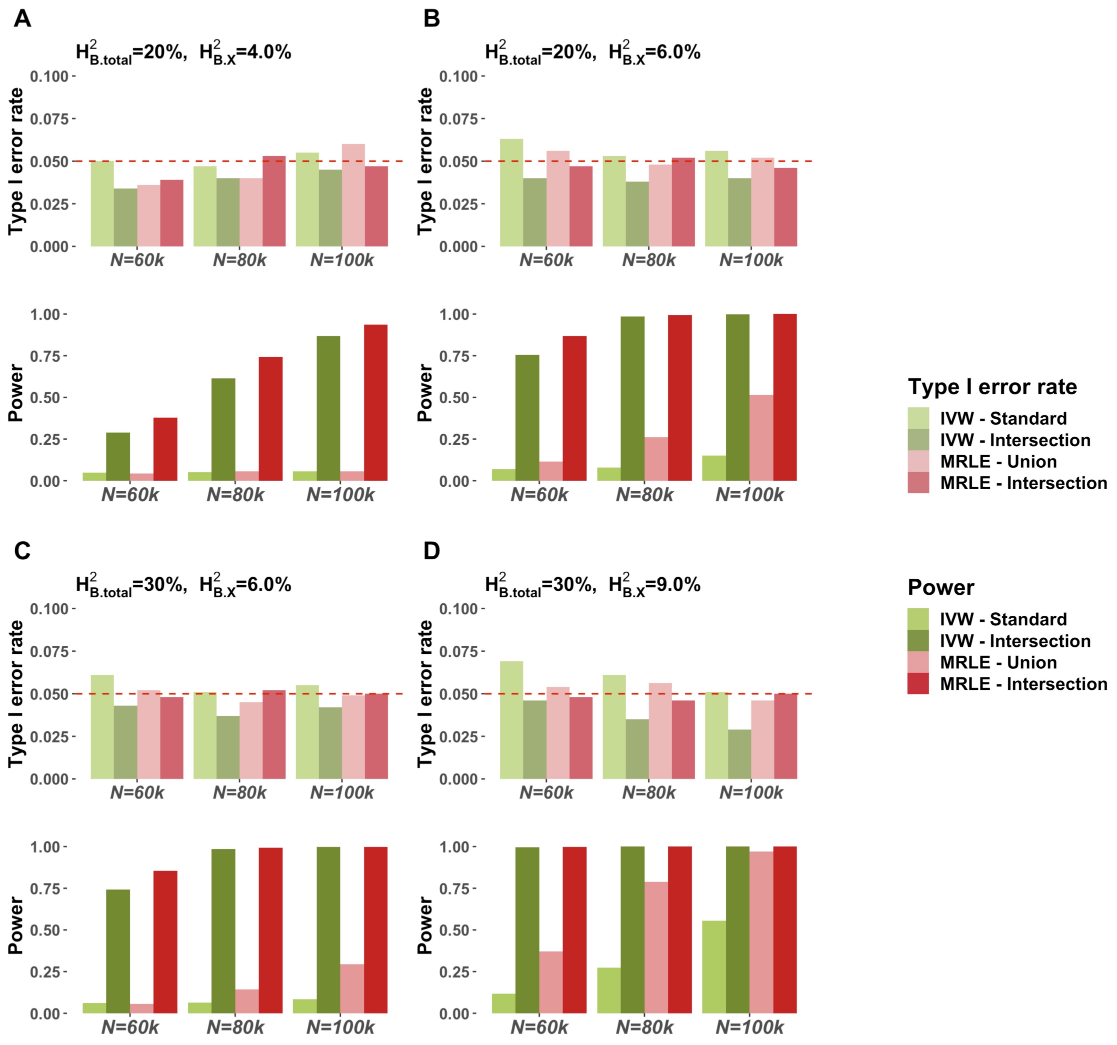


Figure S2. Simulation results assuming a total of $\boldsymbol{K} = 4$ biomarkers based on 1000 simulations per setting. $H^2_{\text{B.total}}$ and $H^2_{\text{B.X}}$ denote the total genetic heritability of each biomarker and the heritability of each biomarker explained by the latent exposure, respectively. IVs are defined as either the SNPs associated with at least one biomarker ("IVW-Standard" and "MRLE-Union", $\alpha = 5 \times 10^{-8}$) or the SNPs associated with at least two biomarkers ("IVW-Intersection" and "MRLE-Intersection", $\alpha = 5 \times 10^{-6}$). In each subfigure, the upper panel shows the empirical type I error rate under $\theta = 0$, and the lower panel shows the empirical power under $\theta = 0.1$.

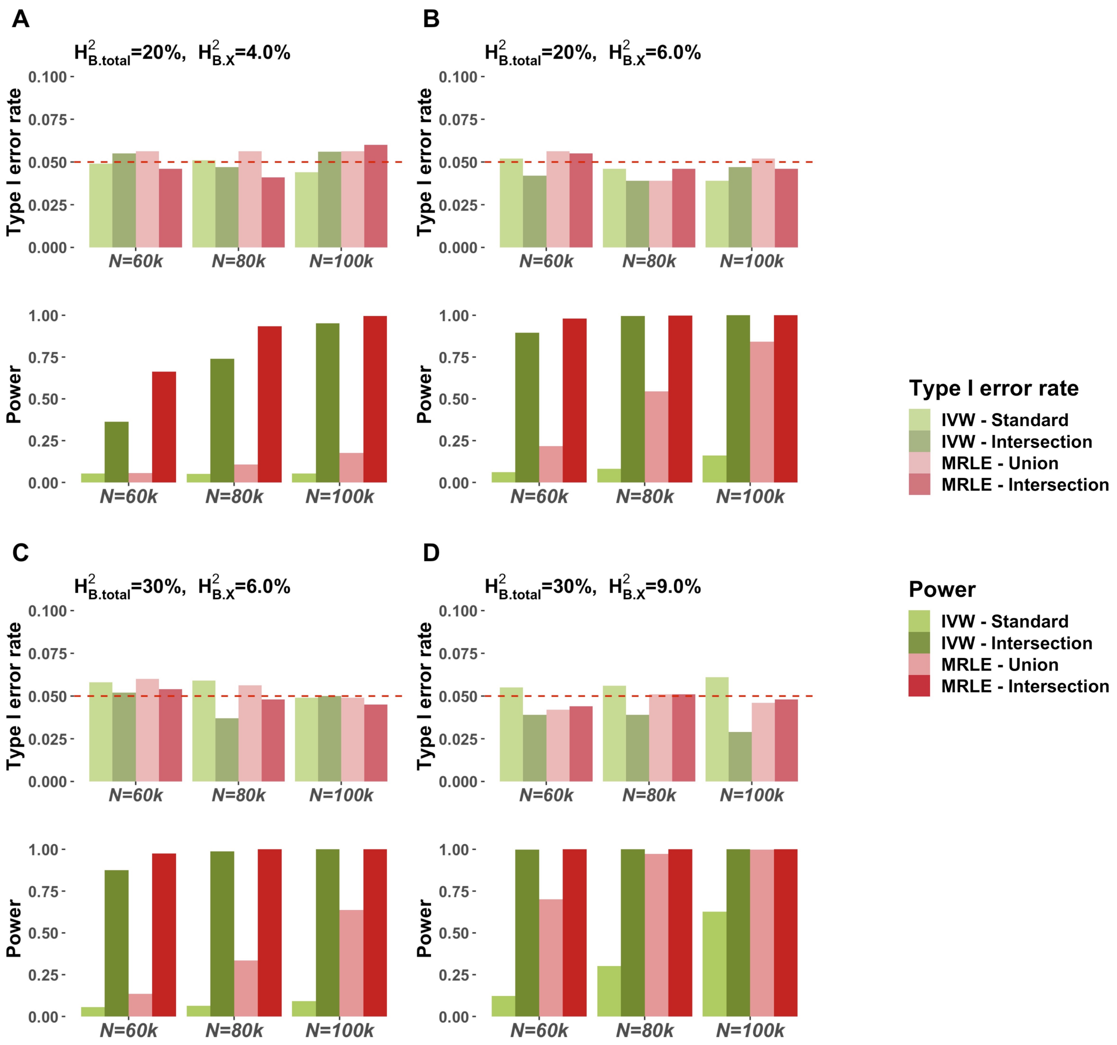


Figure S3. Simulation results assuming a total of $\boldsymbol{K} = \mathbf{8}$ biomarkers based on 1000 simulations per setting. $H^2_{\text{B.total}}$ and $H^2_{\text{B.X}}$ denote the total genetic heritability of each biomarker and the heritability of each biomarker explained by the latent exposure, respectively. IVs are defined as either the SNPs associated with at least one biomarker ("IVW-Standard" and "MRLE-Union", $\alpha = 5 \times 10^{-8}$) or the SNPs associated with at least two biomarkers ("IVW-Intersection" and "MRLE-Intersection", $\alpha = 5 \times 10^{-6}$). In each subfigure, the upper panel shows the empirical type I error rate under $\theta = 0$, and the lower panel shows the empirical power under $\theta = 0.1$.

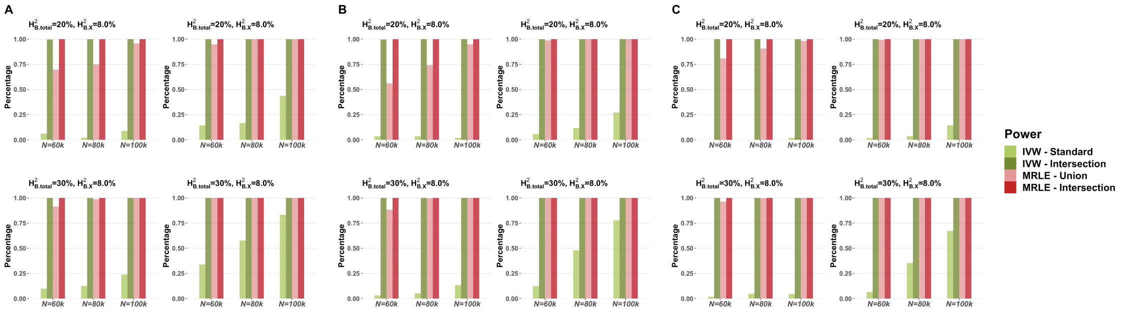


Figure S4. Percentage of rejections that correctly identified the direction of the effect assuming $\theta = 0.1$ and no pleiotropy based on 1000 simulations per setting. $H^2_{\mathrm{B.total}}$ and $H^2_{\mathrm{B.X}}$ denote the total genetic heritability of each biomarker and the heritability of each biomarker explained by the latent exposure, respectively. IVs are defined as either the SNPs associated with at least one biomarker ("IVW-Standard" and "MRLE-Union", $\alpha = 5 \times 10^{-8}$) or the SNPs associated with at least two biomarkers ("IVW-Intersection" and "MRLE-Intersection", $\alpha = 5 \times 10^{-6}$). The number of biomarkers are set to (A) $K = 4$, (B) $K = 6$, or (C) $K = 8$.

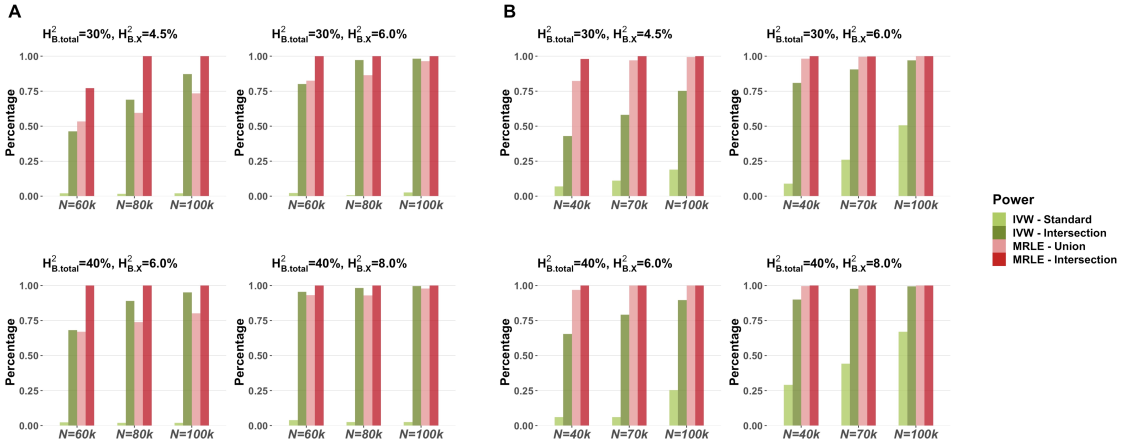


Figure S5. Percentage of rejections that correctly identified the direction of the effect assuming $\theta = 0.1$ and $K = 6$ biomarkers based on 1000 simulations per setting. $H^2_{\text{B.total}}$ and $H^2_{\text{B.X}}$ denote the total genetic heritability of each biomarker and the heritability of each biomarker explained by the latent exposure, respectively. IVs are defined as either the SNPs associated with at least one biomarker ("IVW-Standard" and "MRLE-Union", $\alpha = 5{\times}10^{-8}$) or the SNPs associated with at least two biomarkers ("IVW-Intersection" and "MRLE-Intersection", $\alpha = 5 \times 10^{-6}$). **(A)** Results assuming correlated pleiotropic effects between the biomarkers and the outcome; **(B)** results under correlated pleiotropic effects across biomarkers.

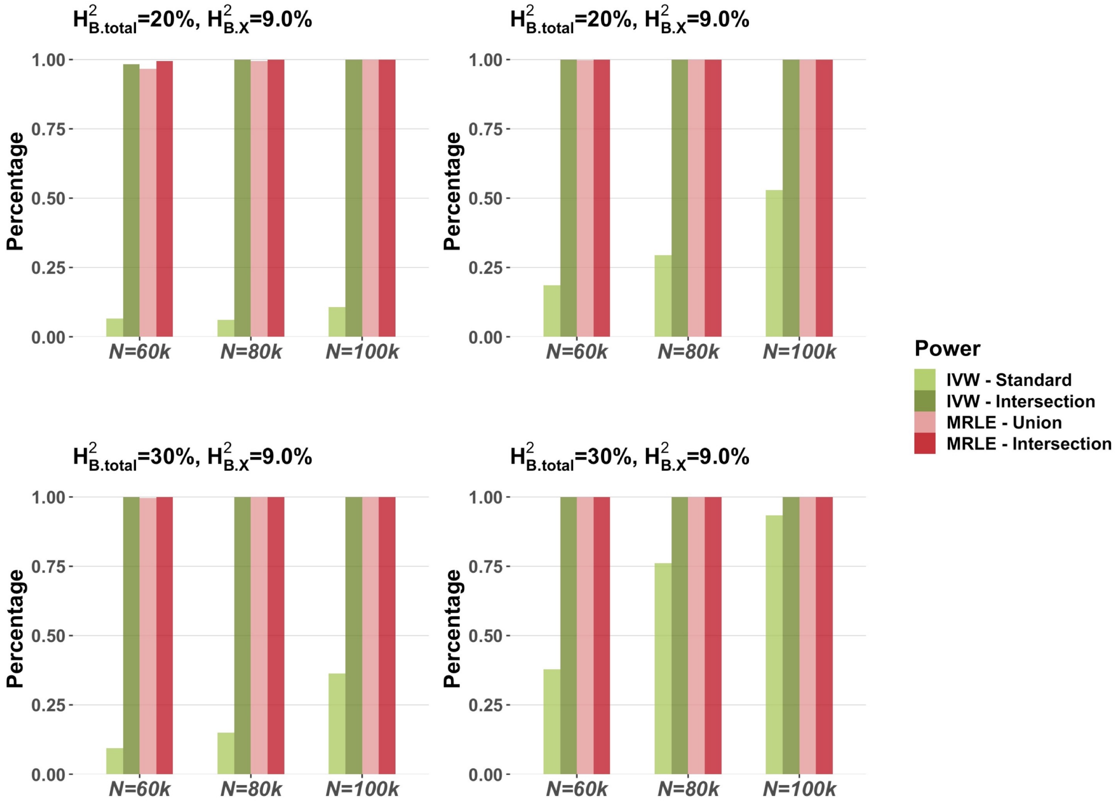


Figure S6. Percentage of rejections that correctly identified the direction of the effect assuming correlated pleiotropic effects across latent exposure and biomarkers given $\theta = 0.1$ and $K = 6$ biomarkers based on 1000 simulations per setting. $H^2_{\text{B.total}}$ and $H^2_{\text{B.X}}$ denote the total genetic heritability of each biomarker and the heritability of each biomarker explained by the latent exposure, respectively. IVs are defined as either the SNPs associated with at least one biomarker ("IVW-Standard" and "MRLE-Union", $\alpha = 5 \times 10^{-8}$) or the SNPs associated with at least two biomarkers ("IVW-Intersection" and "MRLE-Intersection", $\alpha = 5 \times 10^{-6}$).

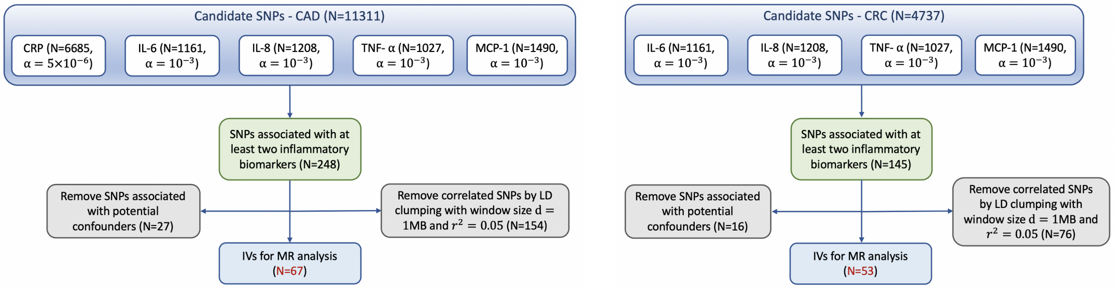


Figure S7: Selection of IVs for the tests on the causal effect of chronic inflammation on CAD (left panel) and CRC (right panel). The selection procedures for RA and PCa are similar to the ones for CAD, and IV selection procedure for EC is similar to the one for CRC. The liberal thresholds for IV selection ($\alpha = 5 \times 10^{-6}$ for CRP and $\alpha = 10^{-3}$ for the other biomarkers) were selected to ensure that at least 10 IVs could be selected across all biomarkers.

Table S1. Data sources for GWAS summary-level association statistics.

| | Study | Ancestry | Sample Size | Source |
|---|---|---|---|---|
| **Inflammatory Biomarkers** | | | | |
| CRP | UK Biobank | European | 320041 | — |
| IL-6 | YFS + FINRISK | European | 8189 | Ahola-Olli et al. (2017)[9] |
| IL-8 | YFS + FINRISK | European | 3526 | Ahola-Olli et al. (2017)[9] |
| TNF-$\alpha$ | YFS + FINRISK | European | 3454 | Ahola-Olli et al. (2017)[9] |
| MCP-1 | YFS + FINRISK | European | 8293 | Ahola-Olli et al. (2017)[9] |
| **Outcome** | | | | |
| CAD | CARDIoGRAM | European | 22233 cases, 64762 controls | Schunkert et al. (2011)[10] |
| RA | Multiple | European + Asian | 29880 cases, 73758 controls | Okada et al. (2014)[11] |
| CRC | UK Biobank | European | 4562 cases, 382756 controls | Zhou et al. (2018)[12] |
| PCa | Multiple | European | 79194 cases, 61112 controls | Schumacher et al. (2018)[13] |
| EC | Multiple (UK Biobank included) | European | 12906 cases, 108979 controls | O'Mara et al. (2018)[14] |

### References


Stephen Burgess, Adam Butterworth, and Simon G Thompson. Mendelian randomization analysis with multiple genetic variants using summarized data. *Genetic epidemiology*, 37(7):658–665, 2013.

Guanghao Qi and Nilanjan Chatterjee. Heritability informed power optimization (hipo) leads to enhanced detection of genetic associations across multiple traits. *PLoS genetics*, 14(10):e1007549, 2018.

Lars Peter Hansen. Large sample properties of generalized method of moments estimators. *Econometrica: Journal of the Econometric Society*, pages 1029–1054, 1982.

Zvi Griliches, Robert Engle, Michael D Intriligator, James J Heckman, Dan McFadden, and Edward E Leamer. *Handbook of econometrics*. Elsevier, 1983.

KW Newey and Daniel McFadden. Large sample estimation and hypothesis. *Handbook of Econometrics, IV, Edited by RF Engle and DL McFadden*, pages 2112–2245, 1994.

Dong-Hui Li, Masao Fukushima, Liqun Qi, and Nobuo Yamashita. Regularized newton methods for convex minimization problems with singular solutions. *Computational optimization and applications*, 28(2):131–147, 2004.

Jin-yan Fan and Ya-xiang Yuan. On the quadratic convergence of the levenberg-marquardt method without nonsingularity assumption. *Computing*, 74(1):23–39, 2005.

Bycroft, Clare and Freeman, Colin and Petkova, Desislava and Band, Gavin and Elliott, Lloyd T and Sharp, Kevin and Motyer, Allan and Vukcevic, Damjan and Delaneau, Olivier and O'Connell, Jared and others. The uk biobank resource with deep phenotyping and genomic data. *Nature*, 562(7726):203–209, 2018.

Ari V Ahola-Olli, Peter Würtz, Aki S Havulinna, Kristiina Aalto, Niina Pitkänen, Terho Lehtimäki, Mika Kähönen, Leo-Pekka Lyytikäinen, Emma Raitoharju, Ilkka Seppälä, et al. Genome-wide association study identifies 27 loci influencing concentrations of circulating cytokines and growth factors. *The American Journal of Human Genetics*, 100(1):40–50, 2017.

Heribert Schunkert, Inke R König, Sekar Kathiresan, Muredach P Reilly, Themistocles L Assimes, Hilma Holm, Michael Preuss, Alexandre FR Stewart, Maja Barbalic, Christian Gieger, et al. Large-scale association analysis identifies 13 new susceptibility loci for coronary artery disease. *Nature genetics*, 43(4):333–338, 2011.

Yukinori Okada, Di Wu, Gosia Trynka, Towfique Raj, Chikashi Terao, Katsunori Ikari, Yuta Kochi, Koichiro Ohmura, Akari Suzuki, Shinji Yoshida, et al. Genetics of rheumatoid arthritis contributes to biology and drug discovery. *Nature*, 506(7488):376–381, 2014.

Wei Zhou, Jonas B Nielsen, Lars G Fritsche, Rounak Dey, Maiken E Gabrielsen, Brooke N Wolford, Jonathon LeFaive, Peter VandeHaar, Sarah A Gagliano, Aliya Gifford, et al. Efficiently controlling for case-control imbalance and sample relatedness in large-scale genetic association studies. *Nature genetics*, 50(9):1335–1341, 2018.

Fredrick R Schumacher, Ali Amin Al Olama, Sonja I Berndt, Sara Benlloch, Mahbubl Ahmed, Edward J Saunders, Tokhir Dadaev, Daniel Leongamornlert, Ezequiel Anokian, Clara Cieza-Borrella, et al. Association analyses of more than 140,000 men identify 63 new prostate cancer susceptibility loci. *Nature genetics*, 50(7):928–936, 2018.

Tracy A O'Mara, Dylan M Glubb, Frederic Amant, Daniela Annibali, Katie Ashton, John Attia, Paul L Auer, Matthias W Beckmann, Amanda Black, Manjeet K Bolla, et al. Identification of nine new susceptibility loci for endometrial cancer. *Nature communications*, 9(1):1–12, 2018.

[*Received February, 2023*]
